# Phenelzine-based probes reveal Secernin-3 is involved in thermal nociception

**DOI:** 10.1101/2023.02.02.526866

**Authors:** Katelyn A. Bustin, Kyosuke Shishikura, Irene Chen, Zongtao Lin, Nate McKnight, Yuxuan Chang, Xie Wang, Jing Jing Li, Eric Arellano, Liming Pei, Paul D. Morton, Ann M. Gregus, Matthew W. Buczynski, Megan L. Matthews

## Abstract

Chemical platforms that facilitate both the identification and elucidation of new areas for therapeutic development are necessary but lacking. Activity-based protein profiling (ABPP) leverages active site-directed chemical probes as target discovery tools that resolve activity from expression and immediately marry the targets identified with lead compounds for drug design. However, this approach has traditionally focused on predictable and intrinsic enzyme functionality. Here, we applied our activity-based proteomics discovery platform to map non-encoded and post-translationally acquired enzyme functionalities (e.g. cofactors) *in vivo* using chemical probes that exploit the nucleophilic hydrazine pharmacophores found in a classic antidepressant drug (e.g. phenelzine, Nardil ^®^). We show the probes are *in vivo* active and can map proteome-wide tissue-specific target engagement of the drug. In addition to engaging targets (flavoenzymes monoamine oxidase A/B) that are associated with the known therapeutic mechanism as well as several other members of the flavoenzyme family, the probes captured the previously discovered *N*-terminal glyoxylyl (Glox) group of Secernin-3 (SCRN3) *in vivo* through a divergent mechanism, indicating this functional feature has biochemical activity in the brain. SCRN3 protein is ubiquitously expressed in the brain, yet gene expression is regulated by inflammatory stimuli. In an inflammatory pain mouse model, behavioral assessment of nociception showed *Scrn3* male knockout mice selectively exhibited impaired thermal nociceptive sensitivity. Our study provides a guided workflow to entangle molecular (off)targets and pharmacological mechanisms for therapeutic development.

## 1. INTRODUCTION

Phenelzine (PHZ, Nardil^®^) is an FDA-approved drug for treating depression (Fiedorowicz and Swartz, 2004). Since it was first introduced into psychiatry more than 70 years ago when tuberculosis patients showed ‘side effects’ of improved mood (Selikoff and Robitzek, 1952), its off-label use has had demonstrated efficacy in a broad range of diseases. For example, PHZ mitigates symptoms of other psychiatric disorders including post-traumatic stress (Zhang and Davidson, 2007), panic and social anxiety (Sheehan et al., 1980; Buigues and Vallejo, 1987; Williams et al., 2020), bipolar (Mallinger et al., 2009), atypical depression (Pande et al., 1996), and treatment-resistant depression, where it outperforms other antidepressants (Kim et al., 2019). PHZ also exhibits analgesic and nociceptive effects in various disease models of pain. For example, anti-nociceptive effects of PHZ were observed in models of inflammatory pain (Mifflin et al., 2016), chronic pain (Davidson and Raft, 1985), in allodynia associated with experimental autoimmune encephalomyelitis (EAE) in a mouse model for multiple sclerosis (Potter et al., 2016) and in a clinical study evaluating PHZ for treatment of neuropathic pain in patients with fibromyalgia (Tort et al., 2012). Further, by acting as an aldehyde scavenger, PHZ exhibits neuroprotective effects in Parkinson’s disease and spinal cord contusive injury (MacKenzie et al., 2010; Song et al., 2010; Al-Nuaimi et al., 2012; Chen et al., 2016; Matveychuk et al., 2022), and reduces oxidative damage following traumatic brain injury (Hill et al., 2020). Importantly, PHZ is not addictive (Ananth et al., 1995) and has anti-abuse potential, as clinical studies revealed PHZ reduces tobacco dependence (George and Weinberger, 2008) and cocaine abuse (Golwyn, 1988). Beyond diseases of the central nervous system (CNS), PHZ was shown to reduce prostate cancer growth and metastasis in preclinical models (Gaur et al., 2019) and attenuate measures of relapse in an open-label Phase II clinical trial in patients with recurrent prostate cancer (Gross et al., 2021). Despite these successes, PHZ has remained a last resort treatment option due to patient tolerability and safety issues including enhanced alcohol toxicity (Weathermon and Crabb, 1999), hypertensive crises (Smookler and Bermudez, 1982), and vitamin B6 deficiency (Malcolm et al., 1994).

PHZ’s antidepressant activity has been attributed to its mechanism of action as irreversible monoamine oxidase A and B inhibitors (MAOIs) (Zeller and Barsky, 1952; Thase et al., 1995), as confirmed by the development of more selective irreversible and reversible next-generation MAOIs for this indication (Gillman, 2011). In a recent study (Lin et al., 2021), our group showed that the promiscuous reactivity of PHZ’s hydrazine group (–NHNH_2_, **Fig. 1A**, highlighted in green) – the brain-penetrant pharmacophore essential for covalent inactivation of MAO via irreversible modification of its flavin adenine dinucleotide (FAD) cofactor by the drug – is likely to facilitate many of the adverse side effects. At the same time, associated off-target effects are also likely to underlie the wide range of off-label non-antidepressant therapeutic actions of this drug reported throughout the literature. Thus, non-MAO protein targets of PHZ represent unexplored therapeutic space for treating neurological disorders. However, mapping target-phenotype relationships and developing selective molecules that modulate their (dys)function in potential disease states requires the development of new tools that can identify and characterize these proteins *in vivo*.

**Figure 1.**
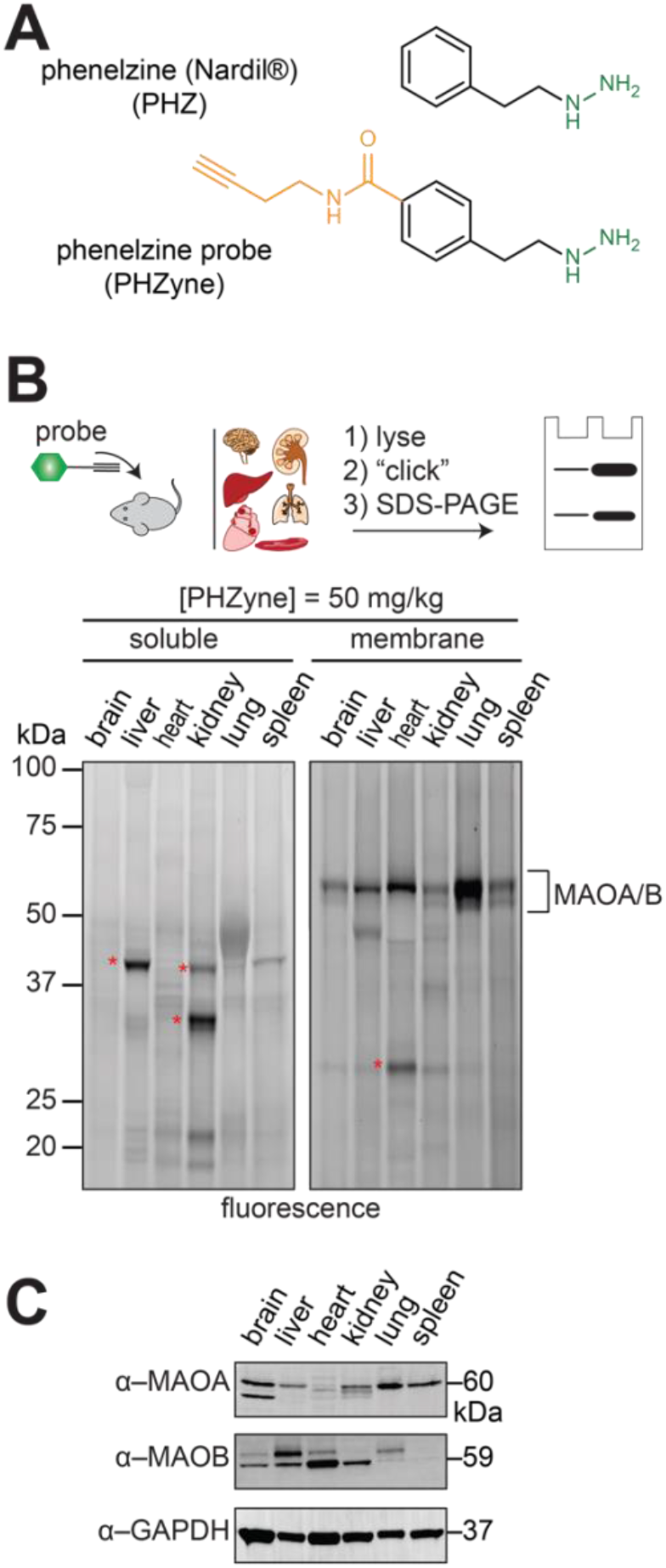
System-wide visualization of proteins covalently modified *in vivo* by PHZyne. **(A),** Structures of phenelzine drug (PHZ) and the corresponding alkynylated probe (PHZyne). The reactive hydrazine pharmacophores are shown in green and chemical modifications made to the parent drug to incorporate the clickable alkyne handle are shown in orange. **(B),** SDS-PAGE analysis of soluble and membrane proteomes from six mouse tissues harvested 4 h post-injection with PHZyne (I.P., 50 mg/kg) and in-gel fluorescence scanning of PHZyne-modified proteins as measured by CuAAC to a rhodamine-azide tag (fluorescent gel shown in grayscale). Florescent protein bands for known on-targets of PHZ are shown by protein name (MAOA/B at ~ 60 kDa); Asterisks (*) highlight unidentified off-targets that remain. **(C)**, Western blot analysis of MAOA/B expression in membrane proteomes across mouse tissues from **(B)** and compared against GAPDH as an approximate loading control. Molecular weights are indicated.

In our prior work, PHZ-based chemical probes were deployed in living cells to capture all MAO and non-MAO hydrazine-reactive proteins in culture (Lin et al., 2021). By conjugating a fluorophore or affinity handle to the alkyne group of the probe (**Fig. 1A**, in orange) using biorthogonal click chemistry, hydrazine-reactive proteins can be visualized by in-gel fluorescence scanning and identified by mass spectrometry-based chemical proteomics (Cravatt et al., 2008; Matthews et al., 2017; Dettling et al., 2020). Using this chemical proteomics discovery platform known as reverse-polarity activity-based protein profiling (RP-ABPP), we demonstrated that the PHZ pharmacophore can functionally inhibit a wide range of non-MAO protein targets from diverse functional classes known to be important in neurological diseases (Lin et al., 2021), but were not previously known to have hydrazine-sensitive activities. Additionally, we established that PHZ-based probes administered to cells can facilitate the discovery of selective small molecules for these proteins despite many considered to be undruggable or have unknown functions. However, in order to evaluate the translational potential of these targets, we sought to expand our approach to be compatible *in vivo*.

In this work, we apply PHZ-based probes as *in vivo* pharmacological tools to discover hydrazine-sensitive biology directly in animal models. We demonstrate that the probes are brain-penetrant following systemic delivery in mice and can be used to measure *in vivo* target engagement and map proteome- and organ-wide targets in the brain and other tissues. In addition, we use activity-based imaging to resolve the spatiotemporal activity of PHZ in cells. To illustrate how the approach can be implemented to deorphanize genes of unknown biological function, we further evaluated Secernin-3 (SCRN3), an uncharacterized metabolic enzyme and unexpected off-target of PHZ in the brain. We show that SCRN3 expression is regulated by pro-inflammatory response pathways induced by lipopolysaccharide (LPS) in macrophages, and that SCRN3 is involved in thermal nociception in male mice. The results illustrate the utility of these drug-inspired probes as *in vivo* pharmacological tools that direct the discovery of genetically-unpredicted protein function in neurological diseases.

## 2. MATERIALS AND METHODS

### 2.1 Animals for target identification *in vivo*

Wild-type C57BL/6J mice (8-14 weeks) were purchased from Jackson Laboratories. Mice were housed with 2-5 mice per cage under a 12-hour normal light-dark cycle (light on AM, light off PM) with ad-libitum access to standard chow and water. Protocols were approved by the Institutional Animal Care and Use Committees (IACUC) at the University of Pennsylvania (Philadelphia, PA), the Children’s Hospital of Philadelphia (Philadelphia, PA).

### 2.2 Clickable probes and drugs

Clickable chemical probes [alkynylated analogs of phenelzine (PHZyne) and phenyhydrazine (PHAyne)] were synthesized and characterized according to previous methods (Matthews et al., 2017; Lin et al., 2021). All probes and drugs [phenelzine (PHZ) and phenylhydrazine (PHA)] were prepared fresh from solid stocks in PBS the day of experiments.

### 2.3 Quantitative chemical proteomic profiling of *in vivo* targets

After being fasted overnight, C57BL/6J male mice (~14 weeks old) were treated (i.p.) with the above probes or drugs (12.5 mg/mL) for 4 h at 50 mg/kg. Mice were euthanized by CO2 asphyxia, followed by cervical dislocation and blood perfusion was performed according to previous methods (Shishikura et al., 2019; Wu et al., 2021). Six major tissues (brain, liver, heart, kidney, lung, and spleen) were harvested, washed with 10 mL ice-cold PBS (3x) and homogenized in PBS or RIPA buffer using a NextAdvance bullet blender. Homogenization beads (~ 50 μL Next Advance, Inc., glass or zirconium oxide beads appropriate for each tissue) were added to each tissue. Following homogenization, (Bullet Blender, BBY24M model, Next Advance, Inc., speed 8, 3 min, 4 °C), samples were diluted in PBS (500 μL) and centrifuged at 1500 x g for 5 min (4 °C) to remove debris. Resultant homogenates were fractionated and prepared as previously described for gel- and MS-based ABPP experiments (Lin et al., 2021), with the exception that 2 mg/mL proteome concentrations were used for experiments.

### 2.4 Gel- and MS-based RP-ABPP

#### Gel-based

To analyze protein activity, soluble and membrane tissue or cell proteomes were conjugated to rhodamine-azide by copper-catalyzed azide-alkyne cycloaddition (CuAAC or ‘click’) chemistry, resolved by SDS-PAGE and visualized by in-gel fluorescence scanning as previously described (Lin et al., 2021).

#### MS-based

Proteomes were similarly conjugated to biotin-azide by CuAAC, probe-labelled proteins were enriched by streptavidin affinity chromatography and digested on-bead with trypsin as previously described (Lin et al., 2021). The resulting tryptic peptide digests from probe-versus drug-treated mice were isotopically differentiated by reductive dimethylation (ReDiMe) of lysine residues using isotopically ‘heavy’ and ‘light’ formaldehyde, respectively (Boersema et al., 2009) as previously described (Matthews et al., 2017), to identify and quantify enrichment and competition heavy/light ReDiMe ratios for PHZyne or PHAyne as defined in Results and **Fig. 3A**.

**Figure 2.**
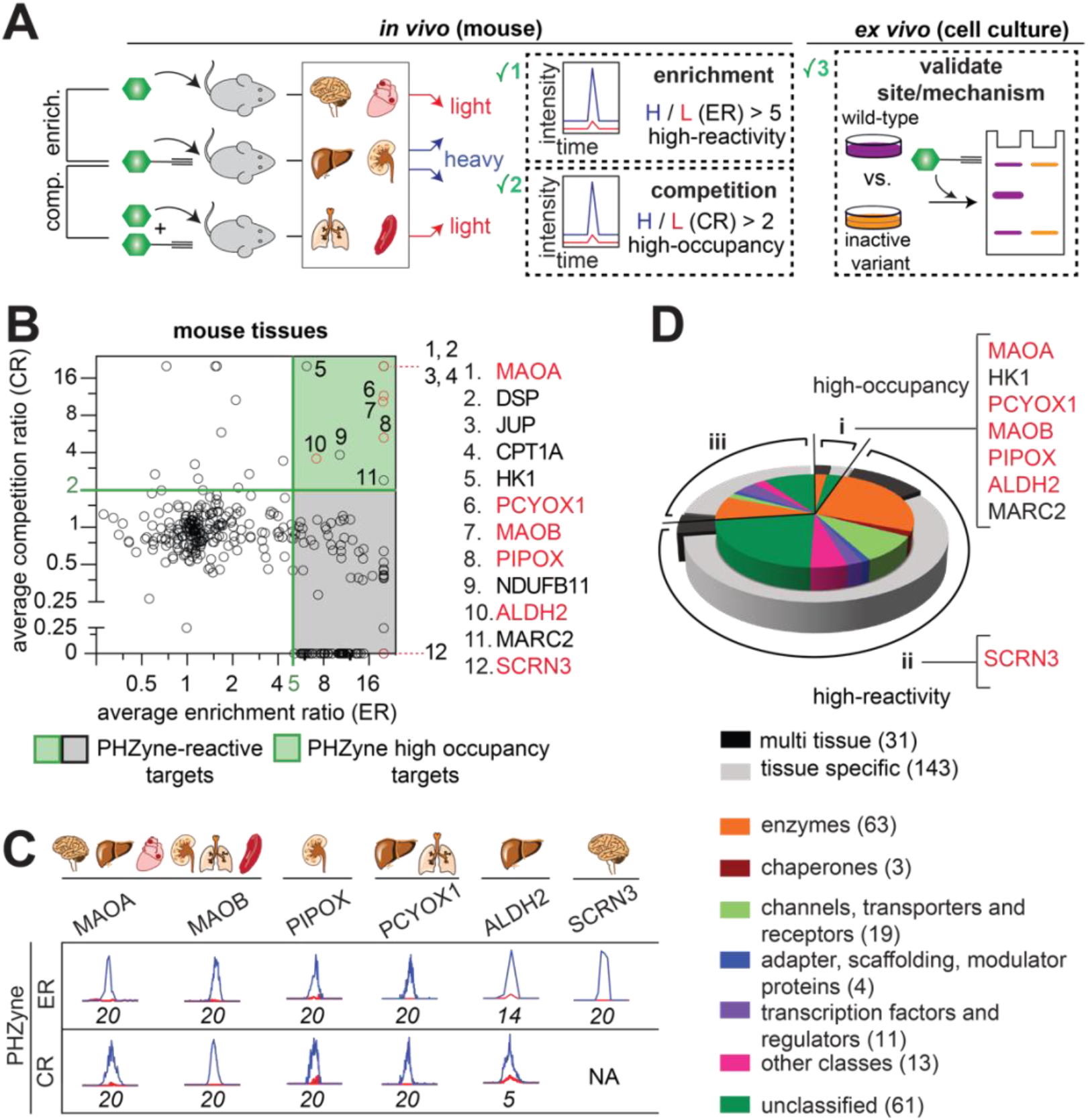
Identification of *in vivo* protein targets of PHZyne in mouse tissues. **(A)**, Schematic for target identification using a probe/drug pair and evaluating the utility of small molecule-protein interactions as described in the text. Proteome- and system-wide quantitative MS-based proteomics experiments (enrichment and competition) identify and quantify probe/drug-target interactions *in vivo* that are reconstituted in cells to validate and characterize the site and mechanistic basis for the interactions captured *in vivo*. **(B),** Quadrant plot of average competition versus enrichment ReDiMe ratios for PHZyne/PHZ targets across all mouse tissues from quantitative proteomic experiments (*n* ≥ 4 / group) in **(A)**. Proteins with ratios that were quantified in enrichment experiments but were either undetected or did not pass the criteria for quantification in corresponding competition experiments were plotted along the x-axis. Proteins with ER ≥ 5 and CR ≥ 2 (upper right quadrant shown in green) were considered both high-reactivity and high-occupancy; listed to the right of the plot. Proteins shown in red are further investigated in this work. **(C)**, Extracted parent ion chromatograms and corresponding heavy/light (H/L) ratios for representative tryptic peptides of PHZyne/PHZ quantified protein targets. **(D)**, PHZyne targets from mouse tissues in groups i-iii (as described in text) classified by tissue-specific reactivity and functional class.

**Figure 3.**
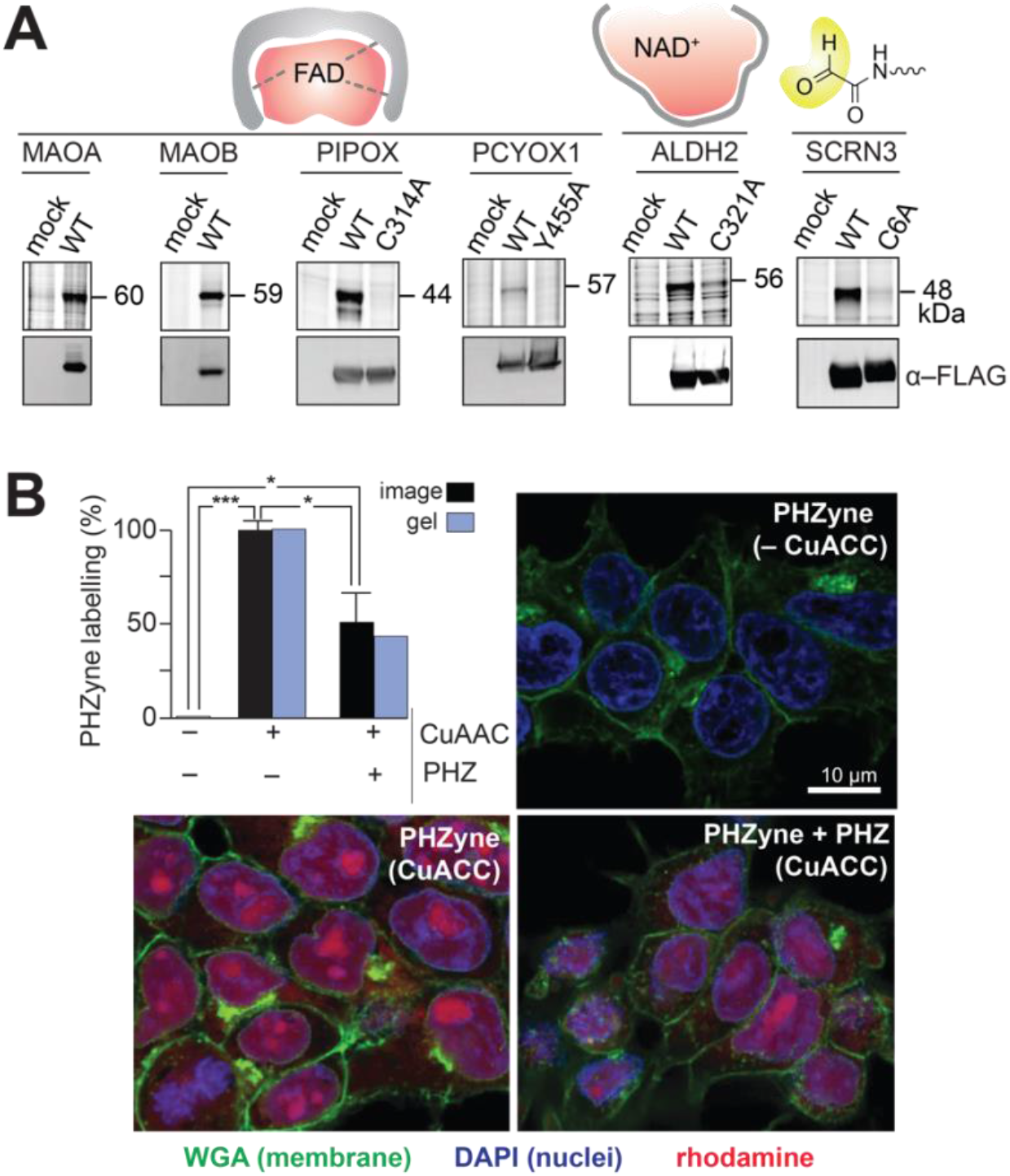
Validation of high-occupancy protein targets in cells. **(A),** Probe labelling (*upper*) and Western blots (*lower*) for PHZyne-treated cells expressing the indicated flavin adenine dinucleotide (FAD)-dependent, nicotinamide adenine dinucleotide- (NAD^+^)-dependent or glyoxylyl (Glox)-harboring protein targets versus inactive variants. The first lane in each gel corresponds to a control transfection (“mock”). Molecular weights are indicated. **(C),** Quantitative image-based analysis of PHZyne/PHZ reactivity from intact cells compared to gel-based analysis of the harvested proteomes (*upper left*). Gel band intensities were calculated as previously described (Lin et al., 2021). PHZyne labelling in the absence (*upper right*) and presence (*bottom left*) of CuAAC and in competition with PHZ (*bottom right*). Rhodamine was conjugated to PHZyne by CuAAC (magenta). Nuclei and membranes were stained with DAPI (blue) and WGA (green), respectively. Mean rhodamine fluorescence intensity was quantified using ImageJ (*n* = 3 per group), and group differences were evaluated by ANOVA followed by Bonferroni Post-hoc analysis (scale bar = 10 μm; * *p* < 0.05, *** *p* < 0.001).

### 2.5 *In vitro* validation of PHZyne and PHAyne mouse targets

#### Gel-based ABPP

Recombinant expression and probe labeling of mouse targets in HEK293T cells were performed as previously described (Lin et al., 2021). Full-length mouse cDNAs encoding MAOA, MAOB, PIPOX, PCYOX1, ALDH2 and SCRN3 were purchased or amplified from an in-house generated cDNA library from Neuro-2a cells (ATCC CCL-131), and subcloned into pRK5-FLAG vectors as described previously (Matthews et al., 2017; Lin et al., 2021). Active site mutants were generated by QuikChange XLII site-directed mutagenesis (Agilent), using primers containing the desired mutations and respective compliments. See **Table S2** for constructs.

#### Western Blotting

Protein expression was analyzed as previously described (Lin et al., 2021). The primary antibodies and dilutions used were as follows: anti-FLAG (1:2500, F1804, Sigma), anti-MAOA (1:1000, PA5-79623, Invitrogen), anti-MAOB (1:1000, PA5-79624, Invitrogen), anti-GAPDH (1:1000, MA515738, Invitrogen), and anti-SCRN3 against His315-Ser814 (1:10,000, NP_083298.1; custom-made rabbit polyclonal antibody from Abclonal, which we verified for specificity in a SCRN3 HEK-293T overexpression system (Matthews, 2017 #571). The secondary antibodies and dilutions used are as follows: goat anti-mouse (1:10,000, ab150113, Abcam), goat anti-rabbit (1:10,000, A-11034, Invitrogen) and goat-anti rabbit HRP (1:2,000, 65-6120, Invitrogen).

#### Site-specific and mechanism-based chemical proteomic experiments

To confirm that Glox is conserved in mouse SCRN3 and captured by PHZyne in the brain *in vivo*, we isolated and characterized the PHZ-modified Glox-containing peptide from probe-treated HEK293T cells expressing recombinant mouse SCRN3 using previously described isoTOP-ABPP methods (Lin et al., 2021) as adapted from previous studies (Weerapana et al., 2007; Matthews et al., 2017).

### 2.6 Quantitative MS-based chemical proteomic profiling

Peptide mixtures from tissue proteomes of treated mice were desalted prior to identification (based on MS2 spectra assignments) and quantification (based on intensity of corresponding extracted MS1 parent ion chromatograms) by liquid chromatography-tandem mass spectrometry-based proteomics (LC-MS/MS), according to previously reported methods (Lin et al., 2021). Proteomic data was processed as previously described (Lin et al., 2021). Protein enrichment ratios (ER) and competition ratios (CR) from soluble and membrane proteome fractions in each tissue were determined from median heavy/light ReDiMe peptide ratios derived from three or more unique quantified peptides per protein and averaged across replicates (*n* ≥ 2) to generate the final ratios reported in **SI Fig. 4**, **Fig. 4E, F**, and **Supplementary Datasets 1** and **2**. Target enrichment and competition were then classified based on ER and CR cut-off criteria, categorized as described in Results.

**Figure 4.**
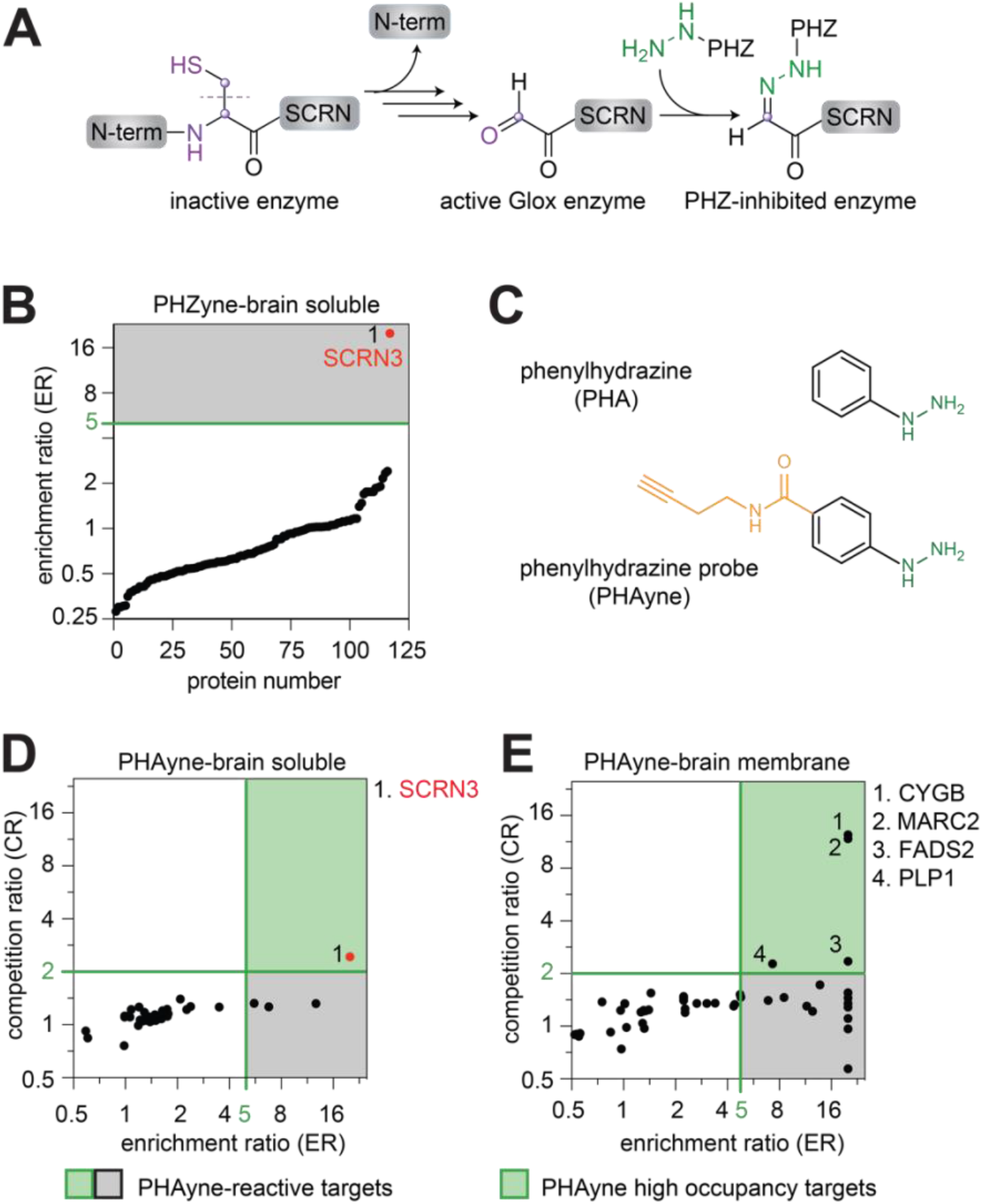
Glox-SCRN3 is an off-target of PHZ in the brain. **(A)**, Schematic for *N*-terminal processing, Glox formation at Cys6 and basis for its reactivity and subsequent capture by PHZ.. **(B)**, Quadrant plot of average enrichment ReDiMe ratios versus protein number for PHZyne from quantitative proteomics experiments (*n* = 11) in the soluble proteomes of mouse brain. Protein targets with ER ≥ 5 (gray box) were considered high-reactivity targets and are annotated. Proteins highlighted in red are investigated in this work**. (C)**, Structures of phenylhydrazine (PHA) and the corresponding alkyne-modified probe (PHAyne) used previously. Green designates hydrazine reactive groups and orange highlights the alkyne handle for detection and identification **(D)**, Quadrant plots of average competition versus enrichment ReDiMe ratios for PHAyne from quantitative proteomic experiments in the soluble (*n* = 2) and **(E)**, membrane (*n* = 2) proteomes of mouse brain. High-occupancy PHAyne targets and listed to the right of the plots. Proteins highlighted in red are investigated in this work.

### 2.7 Activity-based imaging in cells

For confocal fluorescence microscopy imaging experiments, 4 x 10^5^ HEK293T cells were plated onto glass coverslips (P7405, Sigma). The next day, cells were washed and treated with PHZyne/PHZ under identical conditions as gel - and MS-based experiments described previously (Lin et al., 2021). Following treatment, cells were washed and fixed for 20 min with 4% paraformaldehyde (PFA) in PBS (157-4, Electron Microscopy Sciences). For immunofluorescent detection and click chemistry, fixed cells were incubated at room temperature on a shaker, with 2x PBS washes for 10 min between each processing step. Cells were first incubated with WGA-488 (W112961, Thermo) at 5 μg/mL in PBS for 10 min to stain plasma membranes, permeabilized with 0.1% Triton X-100 in PBS for 2 min, conjugated to rhodamine-azide by CuAAC as previously described (Presolski et al., 2011) to visualize PHZyne labelling and incubated with 300 nM DAPI (D9542, Sigma) for 5 min to stain nuclei. For image acquisition, processed coverslips were mounted onto microscope slides with 50 μL of Fluoromount-G (0100-01, Southern Biotech). Images were acquired on a confocal Nikon C2 fluorescence microscope at 20X magnification for quantification and 63X oil for representative images.

### 2.8 Investigation of SCRN3 neurobiology

#### LPS-stimulated RAW 264.7 cells

Murine RAW 264.7 macrophages (ATCC TIB-71) were seeded and cultured similarly to HEK293T cells as described previously (Lin et al., 2021). After 24 h, cells were stimulated with ultrapure lipopolysaccaride (1 μg/mL, LPS, *E. coli* 0111:B4, InvivoGen), harvested and analyzed as described below at 0, 6, 12 and 24 h following LPS treatment.

#### Generation of Scrn3^(-/-)^ mice

*Scrn3* was selected by the NIH International Knockout Mouse Consortium (Brown and Moore, 2012) in 2016. Resulting CRISPR-edited constitutive *Scrn3*^-/-^ knockout mice [strain C57BL/6NJ-Scrn3^em1(IMPC)J/Mmjax^; #042372-JAX] were purchased by Jackson Laboratories. Briefly, four guide RNAs (TAGTGATCGCCTCTTCCCAA, GGAAGAGGCGATCACTATGA, CACTAATTAGTCCAAGCGCA and TTTCCCACTGTAGATGATGT), designed to delete 344 base pairs in exon 3 of the *secernin3* (*Scrn3*) gene, and Cas9 nuclease were introduced into C57BL/6NJ-derived fertilized eggs. Embryos were transferred into pseudo-pregnant females and bred. This mutant results in an amino acid change after residue 53 and early termination 3 amino acid residues later. WT and *Scrn3* knockout C57BL/6NJ mice were generated in Matthews Laboratory from *Scrn3*^(+/-)^ x *Scrn3*^(-/+)^ heterozygous breeding pairs. *Scrn3*^(-/-)^ homozygous KO mice were born at the expected Mendelian frequency and were fertile, thus used to expand the colony. Preliminary characterization of these mice for anatomical and histological abnormalities are still ongoing, but as of yet, the *Scrn3* KOs appear normal.

#### qPCR

mRNA expression levels were measured using real-time quantitative RT-PCR (qPCR). Total RNA was purified from organs with the RNeasy Mini Kit and DNase I (Qiagen)/cDNA was synthesized with iScript™ Reverse Transcriptase (Bio Rad). qPCR was performed with Mx3000P (Agilent) and SYBR^®^ Green PCR Master Mix (Thermo Fisher Scientific). The following primer sequences were used: mouse *Scrn3* forward 5’-CAGTGGCATCAACATGAAGG-3’ and reverse 5’-GCGGAGTTGCAGTAAAGAGG-3’; glyceraldehyde 3-phosphate dehydrogenase (*Gapdh*), forward 5’-TGACAATGAATACGGCTACAGCA-3’ and reverse 5’-CTCCTGTTATTATGGGGGTCTGG-3’. Cycle threshold (Ct) values for organ-wide expression were normalized to total RNA assayed. Relative mRNA levels in RAW264.7 cells were calculated using the ΔΔCt method and normalized to GAPDH mRNA levels (Schmittgen and Livak, 2008).

#### Regional brain tissue collection

Harvested brain tissue (30 g) was sectioned as 1 or 2 mm tissue sections by Rodent Brain Matrix (ASI instruments). The following brain regions were punched by 15 G sample corer (FST) as previously described (Contet et al., 2008; Buczynski et al., 2013):prefrontal cortex (PFC), orbital frontal cortex (OFC), anterior cingulate cortex (ACC) nucleus accumbens (NAcc), dorsal striatum (DS), amygdala (AMYG), lateral hypothalamus (LH), hippocampus (HIPP), thalamus (THAL), ventral tegmental area (VTA), and periaqueductal gray (PAG). Samples were flash frozen in liquid nitrogen and stored at −80°C until analysis. Brain regions were homogenized in PBS by probe sonication and cleared by centrifugation at 200 x g for 2 min (4 °C) for analysis by western blotting.

### 2.9 Evaluation of spinal LPS-induced pain-like behaviors in *Scrn3* KO mice

Male and female WT and *Scrn3* KO C57BL/6NJ mice bred in the Matthews Laboratory were transferred to Virginia Polytechnic Institute and State University (Blacksburg, VA, USA) at 12–16 weeks of age. All behavior protocols were approved by the Institutional Animal Care and Use Committee (IACUC) at Virginia Tech and all experiments complied with the ARRIVE guidelines. The genotype of each genetically altered mouse was confirmed via qPCR. Subjects weighed between 20–30 g and were utilized for testing following exit from quarantine at 16–20 weeks of age.

Intrathecal (IT) injection of LPS was performed according to Hylden and Wilcox (Hylden and Wilcox, 1980). Briefly, mice were anesthetized with isoflurane (2% induction; 1% maintenance) using Somnosuite (Kent Scientific) and a 30G needle affixed to a Hamilton Syringe was inserted percutaneously into the midline between the L5 and L6 vertebrae. Successful entry into the intrathecal space was assessed by the observation of a tail flick. LPS (1μg/5μL PBS) or saline (5 μL PBS) were injected over an interval of 30 s. Mice were removed from anesthetic and allowed to recover in their homecages prior to testing. No motor dysfunction or loss of muscle tone was observed. All experimental procedures were performed during light phase.

Male and female WT and *Scrn3* KO mice were tested for tactile allodynia, changes in grip strength, thermal hyperalgesia, and cold allodynia at different timepoints after IT injection with LPS or saline as follows: tactile at 4, 24 and 48 h; grip force and hotplate at 6, 24 and 48 h and cold at 24 and 48 h. Baselines for each test for each mouse were collected prior to injection procedures and plotted as t = 0 in the time course data. A total of 37 WT (17 male, 20 female) and 34 *Scrn3* KO (16 male, 18 female) mice were used in these studies. Each animal was evaluated in all four tests allowing intermittent resting time between tests to avoid sensitization. All behavioral testing was performed by the same observers who were blinded to the treatment conditions, with animals randomized by another investigator (all female). The observer was unblinded at the conclusion of the experiment.

#### Tactile allodynia

Tactile allodynia was evaluated using manual von Frey filaments with buckling forces between 0.02 and 2 g (Touch Test, Stoelting Co.) applied to the mid-plantar surface of each hind paw using the up-down method as previously described (Chaplan et al., 1994; Gregus et al., 2018). Mice were acclimated to the testing room in a 4-sided acrylic chamber with only one transparent wall (3 x 3 x 7.5 in) and placed on a metal mesh grid under controlled lighting conditions (~100 lux) for a minimum of 60 min (baseline measurements) or 15 min (experimental timepoints). Any mouse exhibiting allodynia prior to IT injection (e.g., basal 50% paw withdrawal threshold (PWT) ≤0.79 g) was excluded from the study. Given that both hind paws are affected by IT injections, thresholds were assessed in both hind paws and averaged for each timepoint.

#### Grip strength

Changes in grip strength were evaluated using a grip force meter (BIOSEB #BIO-GS3) using BIO-CIS response analysis software as previously described (Montilla-Garcia et al., 2017). Briefly, mice were allowed to grasp a metal grid and their tails were gently pulled by hand for 3 s to measure the maximal force exerted by the mouse before releasing the grid. Measurements were made in triplicates to report mean grip force per mouse per time point. Data for each mouse were plotted as mean change in grip force (g) from baseline (prior to IT injection procedures).

#### Thermal hyperalgesia

Thermal hyperalgesia was evaluated by the hot plate assay where mice were tested for withdraw latency as previously described (Naidu et al., 2010). Briefly, mice were habituated to the testing room in their home cages for a minimum of 1 h and acclimated to a hot plate apparatus (IITC, #39) at ambient temperature for a minimum of 10 min before baseline and subsequent test measurements the next day. Thermal paw withdrawal latency at 52.5°C (noxious) was measured in triplicate using a hand timer. Jumping, biting, licking, and clutching of hind paws was considered a nociceptive response as described previously (Jourdan et al., 2001), and a cutoff time of 20 s was used to prevent tissue damage. Data represent the latency (seconds) to respond to the indicated temperature.

#### Cold allodynia

Cold allodynia was evaluated using the cold plantar assay as previously described (Brenner et al., 2012), where withdrawal latency was measured by application of dry ice beneath a glass testing surface. Mice were acclimated to the testing apparatus for 2-3 h until they were calm and paws were flat on the glass testing surface. Given that both hind paws are affected by the IT injection itself, thresholds were assessed in both hind paws and averaged for each timepoint. Data represent the latency (seconds) to respond to the cold stimulus.

#### Data processing and statistical analysis

For each test, time course data for each animal was expressed as hyperalgesic index (% change from baseline x each time point per animal) or area under the curve (AUC) using GraphPad Prism (version 9.4.1) as previously described (Gregus et al., 2018). Statistical analyses were performed using GraphPad Prism. All data are reported as mean ± SEM, and individual data points are indicated. All behavioral tests were analyzed using 2-way ANOVA (sex x genotype) and Bonferroni post hoc. The criteria for significance were as follows: *P<0.05, **P<0.01, ***P<0.001.

## 3. RESULTS

### 3.1 A clickable probe to directly evaluate PHZ-modified proteins *in vivo*

Alkynylated analogs of covalent drugs compatible with late-stage biorthogonal click chemistry developed for ABPP and chemical proteomics approaches has proven powerful for identifying drug targets *in vivo* (Speers et al., 2003; Niphakis et al., 2012; Niphakis and Cravatt, 2014; Zuhl et al., 2016; Niessen et al., 2017; van Esbroeck et al., 2017; Whitby et al., 2017; Caneque et al., 2018). Here, we applied our alkynylated PHZ probe (PHZyne) to directly visualize, identify and quantify reactive and high-occupancy on- and off-targets of PHZ in mouse tissues. We first evaluated the *in vivo* reactivity of PHZyne by systemic PHZ/PHZyne treatment (I.P., 50 mg/kg, 4h), followed by tissue proteome coupling to an azide-rhodamine reporter tag by copper (I)- catalyzed azide-alkyne cycloaddition (CuAAC), SDS-PAGE and in-gel fluorescence scanning (**Fig. 1B**, **Supplementary Figs. 1** and **2**). Consistent with organism-wide covalent inhibition of MAOA/B activity *in vivo*, two PHZyne-labelled protein bands at the expected molecular weight for MAOA/B (~60 kDa) were conserved in membrane proteomes across all six tissues profiled (brain, liver, heart, lung, kidney and spleen). Corresponding protein expression was confirmed by Western blotting (**Fig. 1C**). In addition, multiple off-target protein activities appeared to vary across tissues, suggesting their functions are tissue-specific and PHZ may serve as an *in vivo* active inhibitor for non-MAO targets.

For any *in vivo* active chemical probe (e,g, PHZyne), a specific drug/probe pair (PHZ/PHZyne) must meet three primary criteria to advance a target forward for selective inhibitor development (**Fig. 2A** and **Supplementary Fig. 3**). First, tissue analyzed following systemic administration of the probe (defined by its alkyne handle for biochemical enrichment) must ‘enrich’ for the protein of interest when compared against the native drug (which has no handle for enrichment). If sufficient protein enrichment is achieved from tissues harvested from probe-treated mice versus that of native drug-treated mice (enrichment ratio or ER, Criteria #1), the protein can then be evaluated for ‘competition’ by pretreating animals with the drug prior to treatment with the probe. If the drug blocks enrichment by the probe (competition ratio or CR, Criteria #2), this criteria indicates high reactivity of the target towards the drug and high-occupancy of the target by the drug. This parameter is critical for determining small molecule-target stoichiometry and is useful in assessing the likelihood that engagement of a specific target may produce a therapeutic response (e.g. efficacy) or cause deleterious side effects in patients (e.g. safety). Experimentally, meeting these criteria is essential for downstream structural characterization of previously unknown post-translational modifications, catalytic activities and/or novel molecular mechanisms of enzyme inhibition that were captured by the probes on the protein level *in vivo*. For example, proteins that satisfy Criteria #2 are validated *in vitro* by the presence of a well-defined probe-labeled band that is specific to probe-treated cells that recombinantly express a given target protein. To determine the mechanistic basis for their probe reactivity and to assess the likelihood that a probe/drug-target interaction may contribute to a therapeutic response or deleterious side effect, probe labeling must be lost in cells that express target proteins mutated at the predicted site of probe reactivity that is essential for primary function (e.g. mechanism-based interaction, Criteria #3). Protein targets that meet all three criteria can be selected for inhibitor development by exploiting both the molecular recognition of binding sites for primary substrates and the specific reactivity of a target’s active site. Proteins that fail to meet these criteria require the development of optimized chemical probes tailored for a specific target class or with a specific type of chemical reactivity.

### 3.2 Identification and characterization of PHZyne protein targets in mouse tissues

Using our RP-ABPP chemical proteomics discovery platform (Matthews et al., 2017; Dettling et al., 2020), we identified and quantified protein targets of PHZyne that satisfy Criteria #1 (enrichment by PHZyne) and Criteria #2 (competition by PHZ) in mouse tissues. Protein target enrichment was determined by comparing tissue proteomes from probe-treated (50 mg/kg, IP, 4 h) versus drug-treated (50 mg/kg, IP, 4 h) control mice. Protein target competition was determined by comparing tissue proteomes from probe-treated (50 mg/kg, IP, 4 h) mice in the absence and presence of prior pretreatment with PHZ under the same conditions (50 mg/kg, IP, 4h). PHZyne-labelled targets were then identified from each tissue by CuAAC coupling to a biotin-azide tag, streptavidin enrichment, and quantitative LC-MS using reductive dimethylation (ReDiMe) chemistry to append isotopically “heavy” and “light” methyl groups to tryptic peptides of labelled proteins from probe-treated versus control-treated mice from both types of experiments (**Fig. 2A**). Heavy/light protein enrichment ratios (ER) and competition ratios (CR) were determined by the median ratio of ≥ 3 unique quantified peptides per protein from soluble and membrane proteomes in each tissue (**Supplementary Dataset 1**; *n* ≥ 4/group). To assess global target engagement of PHZ across the whole animal, ratios were averaged across all tissues and corresponding aggregate CRs were plotted against aggregate ERs (**Fig. 2B**). Proteins with ER ≥ 5 across tissues (in green and gray quadrants) were considered PHZyne-reactive, yet many of them were low stoichiometry labeling events (gray quadrant) and thus may not be expected to contribute to a therapeutic effect of PHZ. By contrast, PHZyne-reactive targets that also exhibited aggregate CR ≥ 2 (in green quadrant only), achieve high-occupancy engagement by PHZ *in vivo*, indicating these specific PHZ-target interactions are most likely to be responsible for the therapeutic effects of the drug.

The data suggests that our MAOI-based probe can measure enzyme activity and quantify tissue specific reactivity towards both known (MAOA/B) and previously unknown targets *in vivo* both proteome- and organism-wide. Representative peptide ratios from enrichment and competition experiments for PHZyne-reactive targets, MAOA, MAOB, additional FAD-dependent enzymes, peroxisomal sarcosine oxidase (PIPOX) and prenylcysteine oxidase 1 (PCYOX1), the nicotinamide adenine dinucleotide [NAD(P)^+^]-dependent aldehyde dehydrogenase (ALDH2) and the *N*-terminal glyoxylyl (Glox)-harboring SCRN3, are shown in **Fig. 2C**. MAOA/B are shown to be targets of PHZ across all tissues analyzed. As PHZ is meant to target MAOA/B in the brain as an antidepressant, the lack of tissue specificity of the drug, as well as the MAO activity it inhibits, may contribute to the known adverse effects associated with PHZ in the clinic. ALDH2, responsible for the second step of alcohol breakdown, is another key mitochondrial enzyme involved in monoamine metabolism (Axelrod et al., 1959; Erwin and Deitrich, 1966) that is affected by PHZ, causing patients to experience alcohol toxicity (Weathermon and Crabb, 1999). The known off-target ALDH2 has tissue specific reactivity in the liver, consistent with its role in metabolizing alcohol. Tissue specific reactivity is also observed for the alternative flavoenzymes captured by PHZyne. For example, PIPOX which is expressed in the liver and kidney (Chikayama et al., 2000) exhibited reactivity with PHZyne in the kidney. In contrast, PCYOX1 a lipoprotein-associated protein recently described as a potential target in atherosclerosis (Banfi et al., 2021), exhibited reactivity with PHZyne in liver and lung tissue. Finally, SCRN3 an uncharacterized enzyme and therefore generally rare target exhibited PHZ reactivity in the brain.

Because the representation of the data in **Fig. 2B** serves as an organism-wide view of PHZ target engagement, we generated analogous ratio plots for each tissue individually (**Supplementary Fig. 4**) and performed a meta-analysis to evaluate tissue-specific reactivities. All quantified proteins (658 targets) were categorized into four groups according to their reactivity with PHZyne (ER) and consequent occupancy of the target (CR): i) high-occupancy (ER ≥ 5 *and* CR ≥ 2); ii) high-reactivity undetermined occupancy (ER ≥ 5 *and* CR = unquantified; iii) low reactivity (ER ≥ 5 *and* CR < 2; ER = unquantified *and* CR ≥ 2; ER = 2–5 *and* CR ≥ 2); and iv) nonreactive (ER < 2 *and* CR < 2). Further analysis of classes i–iii (174 targets) predictive of intentional engagement by PHZyne, showed 40% are not found in the DrugBank. This subset of targets was further classified by tissue-specific reactivity and annotated functional class (**Fig. 2D**). Overall, the majority of these proteins exhibited tissue-specific probe reactivity (143, 82%) suggesting their activities are compartmentalized and likely are associated with tissue-specific functions. Additionally, the majority of reactive targets are either known or predicted enzymes (63, 36%) or remain functionally unannotated (61, 35%). Although a minor fraction of targets (50, 29%) represent other functional classes, the data suggests the potential utility of exploiting the reactivity of covalent drug pharmacophores to drive functional annotation of proteins when their activities cannot be readily predicted from gene or protein sequence. As such, our global chemical and biochemical reactivity profiles may serve as a useful tool to help fill the gaps where genetic prediction isn’t working. This is especially true for functionally reactive sites acquired post-translationally as they are installed by dedicated machinery and cellular mechanisms that are not directly genetically encoded into the proteins exploiting them for primary function.

### 3.3 *In vitro* validation of PHZyne protein targets

We first confirmed PHZyne target engagement (Criteria #1 and #2) with the established targets of the drug – MAOA and MAOB (**Fig. 2B**) thereby providing proof-of-concept and feasibility of the approach *in vivo*. We further investigated covalent PHZ inhibition of recombinant mouse MAOA/MAOB, PIPOX, PCYOX1, ALDH2 and SCRN3 proteins using PHZyne-treated HEK293T cells (**Fig. 3A**). Expression of WT mouse proteins furnished strong fluorescent bands at appropriate molecular weights in gel-based probe labelling profiles. Mutating active site residues essential for cofactor binding and catalysis (C314 and Y455 in PIPOX and PCYOX flavoenzymes, respectively and C321 in ALDH2) abolished probe labeling (Criteria #3), confirming the engagement events are active-site directed and mechanism-based, thereby inhibiting function. Finally, mutating the novel Glox-harboring residue that engages PHZ by a divergent mechanism in SCRN3 (C6) (Lin et al., 2021) also abolishes labelling. In summary, the data validate the mechanistic basis for previously unknown PHZ off-targets that result in covalent inhibition of diverse enzyme activities *in vivo*. This insight can be used to improve drug safety by engineering against off-targets (e.g. ALDH2) and additionally uncover unrelated and divergent mechanisms with other targets.

### 3.4 PHZyne for activity-based imaging

Next, we sought to assess whether hydrazine-based ABPP probes may also serve as versatile tools for spatiotemporal mapping by activity-based imaging. For live cell imaging, PHZyne-treated HEK293T cells were visualized by confocal fluorescence microscopy under the same conditions described for the gel- and MS-based proteome profiling experiments (**Fig. 3B** and **Supplementary Fig. 5).** Imaging of PHZyne-sensitive activites was dependent on fluorophore conjugation by CuAAC (upper right versus lower left). Pretreatment with PHZ (lower right) diminished the mean fluorescence intensity by 48% (black bars), consistent with the average extent of competition seen in probe labelling profiles of soluble and membrane proteomes (45%, blue bars) (Lin et al., 2021). These initial data show that probe labelling can be visualized in intact living cells, which we expect will enable both spatially and temporally resolved activity maps of more selective drug-like compounds in intact tissues in the future. Notably, visualization of covalent drug-target interactions with cellular resolution in intact brain tissue has only recently achieved (Pang et al., 2022) using a number of alkynylated CNS drug analogs that had previously been advanced by classical ABPP approaches – among them a covalent MAOI that operates by a slightly different mechanism from that of PHZ.

### 3.5 Investigating the novel phenelzine target SCRN3

Our findings validate the use of PHZyne in mice for the study of MAOA/MAOB and at least three additional non-MAO targets (**Fig. 2C, 3A**) that passed all three selection criteria (**Fig. 2A** and **Supplementary Fig. 3**): i) enrichment by PHZyne probe *in vivo;* ii) competition by PHZ drug *in vivo;* and iii) via a mechanism-based interaction (e.g. the absence of the probe-labeled band for a catalytically inactive mutant enzyme) that led to enrichment and competition by inhibiting primary protein function in the animal. However, a large number of remaining probe-sensitive targets cannot be evaluated by PHZyne alone (**Fig. 2B**, gray box). For example, many that are engaged by PHZyne (enrichment ratios ≥ 5) were not detected in competition experiments due to low protein abundance and/or low occupancy of targets by the drug. However, these reactive targets may still contribute to the wide range of therapeutic effects of PHZ. For example, it is not known how PHZ inhibition of MAOA/B contributes to anti-nociception, and the likelihood that this effect arises from PHZ engagement with a non-MAO target is high according to our profiling data both *in vitro* and *in vivo* (Matthews et al., 2017; Lin et al., 2021). Prior to this work, we demonstrated that Secernin (SCRN) proteins, specifically SCRN2 and SCRN3, are non-MAO PHZyne targets in cells that covalently react with PHZ via their conserved cysteine-derived *N*-terminal Glox groups naturally harbored by these proteins (**Fig. 4A**). Here, SCRN3 was identified as a brain-specific target of PHZyne in mice (**Fig. 2B** gray box, **Fig. 2D** class ii), with the highest measurable enrichment ratio in the brain (Criteria #1, **Fig. 4B**). Structural characterization of the Glox cofactor in mouse SCRN3 was demonstrated by isoTOP-ABPP (**Supplementary Figs. 6** and **7**). PHZyne labelling was abolished by mutation of the Glox-derived cysteine residue in mutant-expressing cells (Criteria #3, **Fig. 3A**), both conferring Glox functionality and its covalent inhibition by PHZ. The data shows that SCRN3’s Glox-dependent activity is conserved in mice, is present in the brain and its inhibition by PHZ might contribute to one or more of the reported therapeutic effects of the drug.

Because PHZyne/PHZ did not fit Criteria #2 for SCRN3, we evaluated a probe that is structurally similar to PHZyne [alkynylated or free phenylhydrazine (PHAyne/PHA, respectively), **Fig. 4C**], that was previously used to discover and characterize the novel Glox group of SCRNs from cells (Matthews et al., 2017). Consistent with our previously reported cell-based reactivity profiles, SCRN3, but not MAOA/B, was a high occupancy target of PHAyne/PHA in the brain when administered to mice *in vivo* under the same conditions as PHZyne/PHZ (Criteria #1 and #2, **Fig. 4D,E**). Covalent modification of Glox by PHAyne was confirmed for mouse SCRN3 (Criteria #3, **Supplementary Figs. 8** and **9**). Collectively, these data show that we furnished *in vivo* active chemical tools to evaluate Glox-SCRN3 function and its inhibition independent of MAO.

### 3.6 Evaluation of SCRN3 neurobiology in mice

To gain insight into function, we reasoned that *Scrn3* expression might be regulated by one or more transcription factors (TFs), and that the identity or identities of these factors might afford clues as the pathway(s) in which the protein participates. Based on enrichment analysis from the chromatin IP (ChIP) Atlas database (http://chip-atlas.org) (Oki et al., 2018), TFs most likely to be associated with *Scrn3* respond to inflammatory stimuli, suggesting a possible role in inflammation. Among them is runt-related transcription factor 1 (RUNX1), which is required for thermal and neuropathic pain (Chen et al., 2006) and regulated by the proinflammatory response ligand lipopolysaccharide (LPS) (Luo et al., 2016; Bellissimo et al., 2020; Huang et al., 2020). As such, we showed that transcript levels of *Scrn3* in macrophages (RAW264.7 cells) are also LPS-sensitive (**Fig. 5A**), hinting that the protein may have a functional role in inflammatory pain pathways and may contribute to the analgesic effect of the drug.

**Figure 5.**
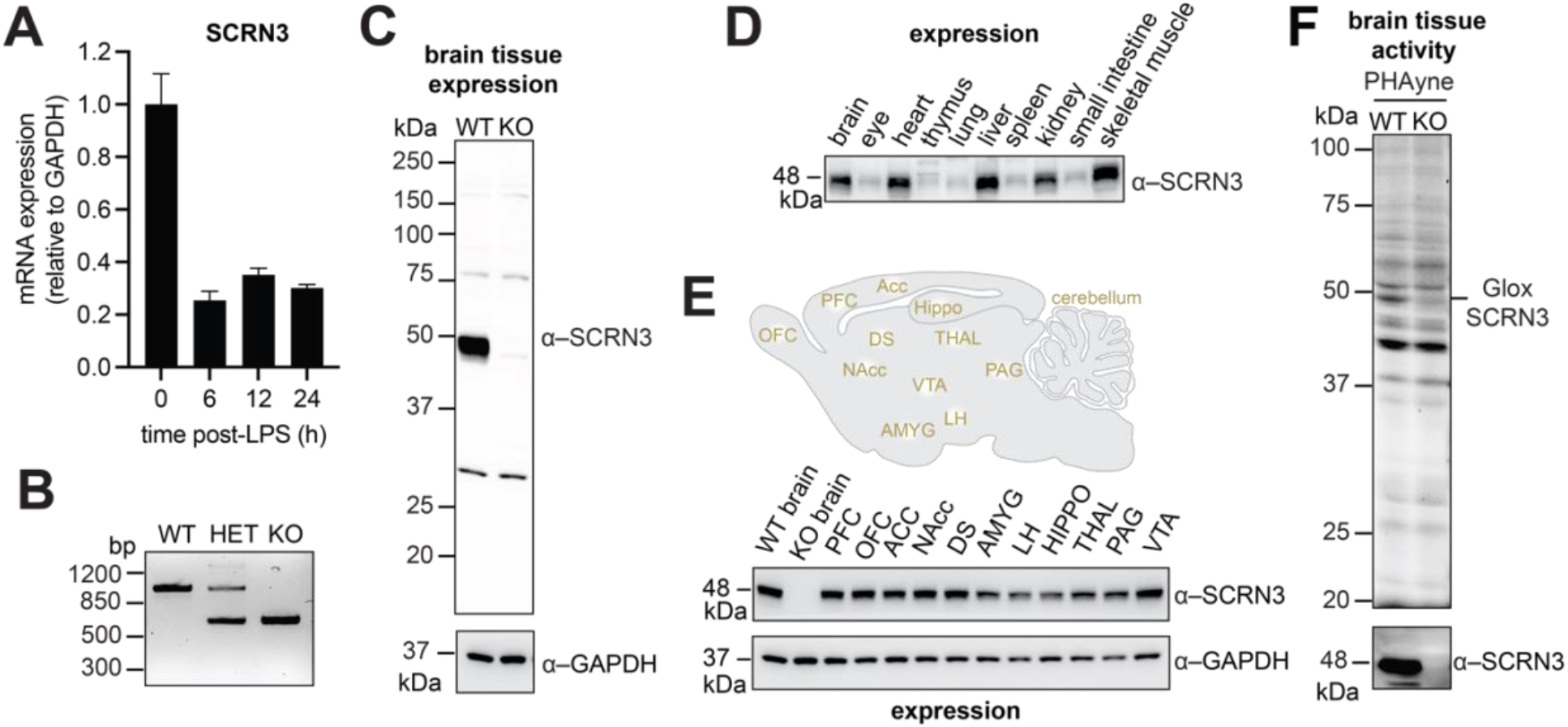
SCRN3 expression and activity in mouse tissues from WT and KO mice. **(A)**, qPCR analysis of *Scrn3* gene expression in macrophages post LPS treatment. Expression was normalized relative to GAPDH. **(B)**, qPCR analysis of *Scrn3* gene expression in WT, SCRN3^(-/+)^ heterozygous, and *Scrn3* ^(+/+)^ homozygous KO mice. *Scrn3* WT and *Scrn3* homozygous KO characteristic bands are around 965 bp and 670 bp, respectively. **(C)**, Western blot analysis of SCRN3 protein expression in mouse brain tissue from WT and *Scrn3* KO mice validating SCRN3-specific antibody. Expression levels were compared to loading control (GAPDH). **(D)**, Western blot analysis of SCRN3 protein expression in ten mouse tissues from WT mice. Coomassie staining was performed as a loading control (See **Supplementary Fig. 10**). **(E),** Western blot analysis of SCRN3 protein expression in mouse brain tissues from WT mice, including regional brain tissues (depicted in brain above), and *Scrn3* KO mice. Expression levels were compared to loading control (GAPDH). Brain tissues from WT and *Scrn3* KO are the same as in **(C). (F)**, *In vivo* reactivity profiles for PHAyne (I.P., 50 mg/kg). in the soluble proteome of mouse brain tissue from WT and *Scrn3* KO mice (*upper*). Western blot analysis of SCRN3 protein expression is below. Molecular weights are indicated.

#### Complementary genetic tools confirm Glox-SCRN3 in the brain

Glox is a conserved functional feature of both human and mouse SCRN3 proteins and their genes share 84% nucleotide sequence similarity. Therefore, we reasoned that a constitutive *Scrn3* KO mouse model could be used to explore the functional significance of mammalian SCRN3 and its druggable Glox group in neurobiology. The discovery of Glox, which was impossible to predict from gene or protein sequence and harbored by proteins of unknown function (SCRN2 and SCRN3 but importantly, not SCRN1), made these genes ideal candidates for NIH’s Knockout Mouse Phenotyping Program (KOMP^2^) (Amos-Landgraf et al., 2022). CRISPR-edited homozygous *Scrn3* KO mice were generated and found to be viable and fertile. They did not exhibit any obvious physiological deficiencies to date, suggesting that deletion of the gene is not critical for growth and development. Absent expression of SCRN3 protein was confirmed by gene sequencing (**Fig. 5B**) and immunoblot analysis (**Fig. 5C**) of KO brain tissue homogenates. Our antibody produced a major band at the predicted molecular weight of full-length SCRN3 protein (48 kDa), that was absent from brain tissue collected from *Scrn3* KO mice (**Fig. 5C**). Contrast to the brain-specific activity observed (**Fig. 2C** and **Fig. 4B**), SCRN3 has comparable tissue expression in the brain, heart, liver, kidney and skeletal muscle (**Fig. 5D** and **Supplementary Fig. 10**). Additionally, regional dissection of brain tissue found that SCRN3 is ubiquitously expressed throughout the CNS and is detectable across multiple corticolimbic brain structures (**Fig. 5E**). Finally, we used the KO animal to confirm the presence of Glox-SCRN3 in the brain of WT mice by *in vivo* capture and labeling by PHAyne (at 48 kDa) that is lost in the KO treated mice (**Fig. 5F**). These probe-labeling profiles reveal that Glox-SCRN3 is present, physiologically active and can be captured with drug-like small molecules in the brain, also highlighting the ability to resolve protein activity from protein expression *in vivo*, a hallmark feature of ABPP. This data also sets the stage to simultaneously evaluate genetic knockdown and pharmacological inhibition of SCRN3 across tissues to probe biochemical and phenotypic function.

#### SCRN3 alters thermal nociception in male mice during a spinal TLR4-dependent, neuropathic-like pain state

Prior to this work, the physiological role of SCRN3 in mammalian biology was unknown. However, in *Drosophila*, a genome-wide screen identified CG10098 (the ortholog of *Scrn3*) as a candidate orphan pain gene that, when deleted, impairs thermal nociception (Neely et al., 2010), suggesting a potential role for SCRN3 in nociceptive processing. The *Drosophila* ortholog is also associated with altered olfactory behavior (He et al., 2016), which indicates the importance of SCRN3 to nervous system function. Additionally, our work (**Fig. 5A**) shows that SCRN3 expression was regulated by Toll-like receptor 4 (TLR4)-dependent pro-inflammatory signaling in mouse macrophages. To evaluate the neurobiological effects of constitutive *Scrn3* gene deletion in mice and to test the hypothesis that SCRN3 has a functional role in thermal nociception, we selected a model for the acute-to-chronic transition of neuropathic pain that is dependent upon the TLR4 receptor (Christianson et al., 2011; Gregus et al., 2018).

Pain, as defined by the International Association for the Study of Pain (IASP), is an unpleasant sensory and emotional experience associated with, or resembling that associated with, actual or potential tissue damage (Raja et al., 2020). Pain processing involves the gating, relay, and interpretation of sensory information transmitted between peripheral tissues and nociceptive neurons and other cell types in ganglia, spinal cord and brain, and thus chronic pain syndromes can be initiated or maintained at peripheral and/or central sites (Basbaum et al., 2009). To evaluate the role of SCRN3 in nociception, four behavioral outputs were measured in both sexes of WT and *Scrn3* KO mice over a 48 h time period following spinal (intrathecal, IT) injection with vehicle or LPS: i) thermal hyperalgesia by the hot plate assay; ii) tactile allodynia by the von Frey test; iii) cold allodynia through the cold plantar assay; and iv) grip force (a functional rheumatological measurement also used in the clinic) using a digital grip force meter. Pain is characterized by allodynia (wherein mechanical, thermal, or chemical stimuli that do not normally produce pain are perceived as painful) or hyperalgesia (a state of enhanced sensitivity to noxious stimuli that is often coupled with spontaneous, non-evoked pain). Grip force is an indicator of muscle strength and nociception, as it is responsive to analgesics (Kehl et al., 2004; Gregus et al., 2021).

Prior to IT injections, baseline measurements for each test showed no statistically significant differences between WT and *Scrn3* KO males and females (**Supplementary Fig. 11**). However, for male mice, two-way repeated measures analysis of variance (ANOVA) revealed a significant effect of genotype and LPS-treatment in two out of four tests (**Fig. 6**). For example, IT delivery of LPS induced a predicted thermal hypoalgesia in WT males (Christianson et al., 2010), whereas this nociceptive phenotype was lost in *Scrn3* KO (**Fig. 6A**). By contrast, there was no effect of IT LPS on grip strength in WT males, however a deficit phenotype (*i.e*., decreased grip strength) was observed in *Scrn3* KO (**Fig. 6D**). As expected, female mice failed to demonstrate genotype- or IT LPS treatment-dependent changes in either behavioral test (**Fig. 6E** and **6H**), consistent with sex-dependent differences in response to TLR4 activation reported in C57BL6J strain (Sorge et al., 2011; Gregus et al., 2021). Both WT and *Scrn3* KO exhibited LPS treatment-dependent increases in tactile allodynia that were conserved in both sexes (**Fig. 6B** and **6F**). Finally, no significant genotype- or treatment-dependent differences were observed in paw withdrawal thresholds from a noxious cold stimulus for either sex (**Fig. 6C** and **6G**). Of note, we recognize that effects on nociception observed in our animals may be impacted by the mixed NJ strain background, as B6N and B6J mice are known to exhibit divergent behavior in some pain models (Ulker et al., 2020). Nonetheless, these findings show that *Scrn3* KO mice are selectively deficient in thermal nociception consistent with that observed in the *Drosophila* screen, indicating that the *Scrn3* gene has a conserved functional role across species.

**Figure 6.**
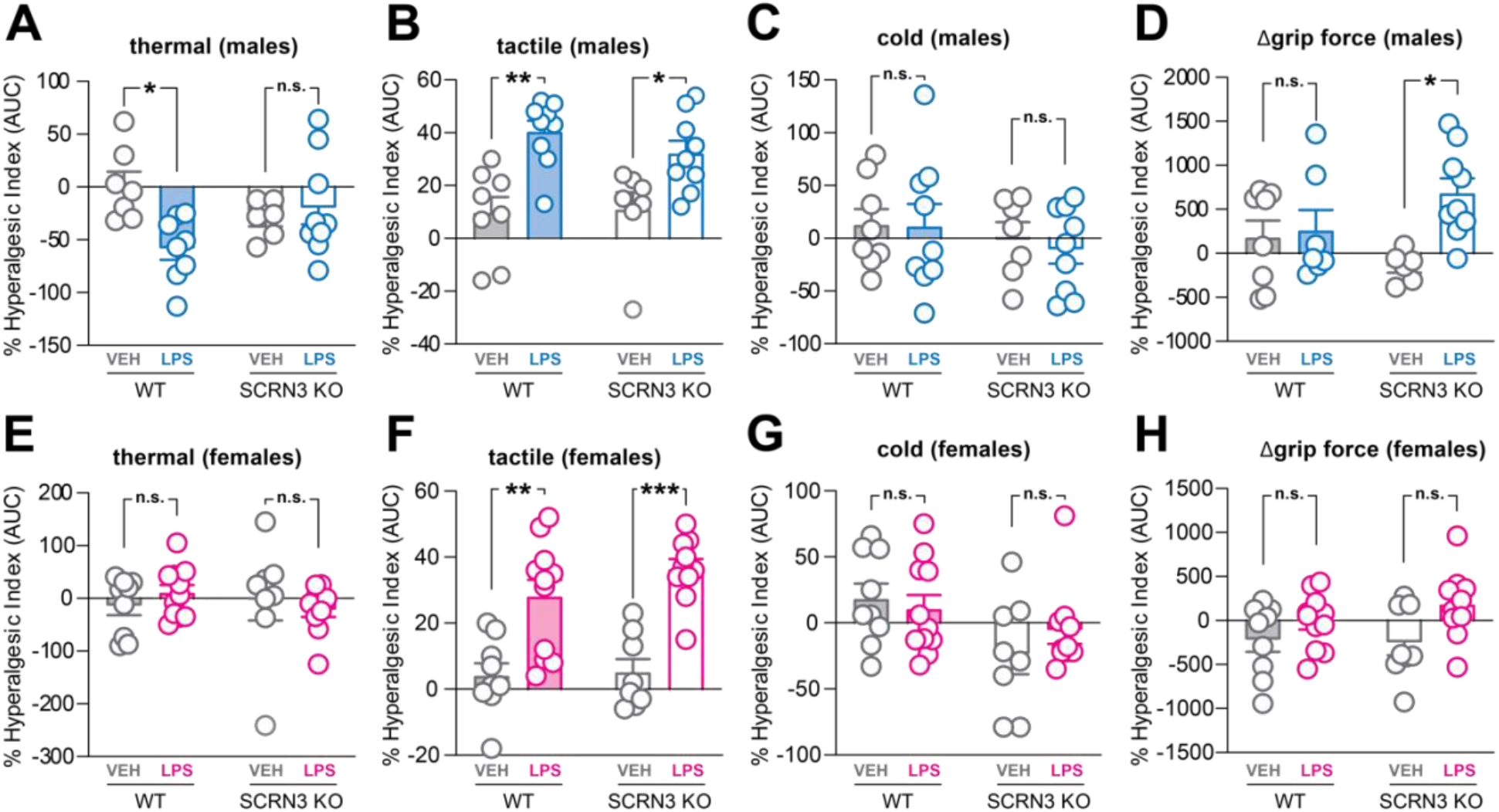
Role of SCRN3 in IT-LPS model of nociception. Male and female WT and *Scrn3* KO mice were treated with 5 μL of vehicle (VEH, saline) or LPS) 1 μg via intrathecal (IT) spinal injection. Behavioral assessment of pain was measured by **(A,E)** thermal hyperalgesia using hotplate assay, **(B,F)** tactile allodynia using von Frey test, **(C,G)** cold allodynia using cold plantar assay, and **(D,H)** paw grip strength deficit using digital grip force meter. All timepoints were used to generate an area under the curve (AUC) assessment, where positive values indicate an increased pain-like response. Data are presented as mean ± SEM, with statistical significance indicated by *P < 0.05, **P < 0.01, ***P < 0.001.

## DISCUSSION

Though MAOA/B is an established drug target of PHZ in the brain, we detected tissue wide expression and activity of MAOA/B with PHZyne, suggesting MAOA/B is physiologically functional in all tissues and supporting the repurposing efficacy of PHZ for other indications. However, it is unlikely that one target is responsible for the many disease indications of PHZ (e.g. analgesics) and our data suggests PHZ’s polypharmacology in other disease states can be evaluated by organ-wide RP-ABPP as well as high-resolution live cell imaging demonstrated here with PHZ-derived chemical probes. If one target, however, is responsible for the therapeutic reach of PHZ, our approach is central to developing tissue-specific targeting for each indication. With the many potentially valuable therapeutic properties, this approach can also be used to develop safer more effective MAOIs that eliminate the effects from unintended off-targets that suboptimally engage with PHZ (e.g. alcohol toxicity via inhibition of ALDH2 activity).

While not all of the enriched targets of PHZyne qualified for selective inhibitor development, they did represent potentially responsive targets for other future probe/drug pairs. For example, we showed that the PHAyne derivative is selective for SCRN3 in the brain and an *in vivo* active compound to further investigate the biological function of SCRN3 through perturbation of its activity and subsequent metabolic pathway dysregulation. Towards this end, we interrogated the neurobiology of SCRN3 in inflammation in macrophages and in nociception using a constitutive *Scrn3* KO mouse model. The relationship between SCRN3 and nociception was predicted in Drosophila, yet it remains unclear the mechanisms by which SCRN3 is correlated to pain processing in higher organisms. In addition, we could not confirm which pharmacological target(s) of PHZ underlies the anecdotal evidence of its efficacy in pain reduction observed in the clinic. Nonetheless, this study demonstrated that SCRN3 is selectively involved in regulating thermal nociception and in grip force deficit (but not in tactile or cold allodynia) following spinal TLR4 activation with LPS, which models TLR4-dependent neuropathic pain chronification. While these data are not conclusive regarding the biological function(s) and therapeutic potential of targeting SCRN3 (or the use of PHZ as a viable therapeutic for chronic pain), they do indicate that *scrn3* gene function is conserved across species, thus enabling future phenotypic studies of SCRN2/3 biology. In addition, this work serves as the basis for establishing chemical and genetic tools to discover the biochemical, metabolic and physiological function of this novel pharmacological target of PHZ. Understanding Glox-SCRN3 function in the brain will heavily rely on its localization, including the specific brain regions, cell types, and neural circuits. Therefore, expanding our *in vitro* imaging data to include recently developed CATCH methods (Pang et al., 2022), that enable brain-wide subcellular imaging of drug-target engagement, will be invaluable for the spatiotemporal resolution of Glox-SCRN3 activity. In addition, coupling these tools to global metabolomics approaches are expected to reveal metabolic activities(s) of Glox-SCRN3.

## CONCLUSION

This work demonstrates how *in vivo* ABPP-based functional proteomics platforms allow for evaluation of drug-target engagement and selectivity across tissues. Moving forward this workflow, illustrated by divergent targets and mechanisms of PHZ inhibition (MAOA/B and SCRN3), provides a roadmap to discover new targets of old drugs *de novo*, supporting the advancement of small molecules defined by and optimized for specific target-phenotype relationships.

## Data Availability

The original contributions presented in this study are included in the supplementary material, further inquiries can be directed to.

## Supplementary Material

Table S1-2 and Supporting figures S1-11 are available as Supplementary Material Supplementary Data Set 1. Compiled PHZyne mouse tissue proteomics data Supplementary Data Set 2. Compiled PHAyne brain tissue proteomics data

## Credit authorship contribution statement

KAB, KS, IC, ZL, PDM, AMG, MWB and MLM designed experiments, interpreted results, and wrote the manuscript. KAB, KS, IC, ZL, NM, YC, XW, JLJ, EA, PDM, AMG and MWB performed investigations. KAB, KS, IC, ZL, AMG, PDM, MWB, and MLM conducted formal analysis. KAB, ZL, AMG, MWB and MLM prepared figures. LP, KS, PDM, AMG, MWB and MLM provided resources and supervision. All authors reviewed and edited the final manuscript.

## Acknowledgements

The authors thank Rick and Janice Woychik and Steve Murray for interest in and generation of the SCRN KO animals (NIH and Jackson Laboratories), synthesis support from Will Parsons (Oberlin College) and computational support from Tammer Ibrahim (University of Pennsylvania), Xian Han (St. Jude Children’s Research Hospital), Robin Park (Integrated Proteomics Applications, Inc), Lin He (Zenagem, LLC) and Radu Suciu (The Scripps Research Institute). We thank Erika J. Olson and Philip E. Dawson (The Scripps Research Institute) for assistance with preparation of the TEV tags.

