## Supplemental Material for "Phenelzine-based probes reveal Secernin-3 is involved in thermal nociception"

### Table S1. Statistics of monoisotopic masses in MS1 spectra for chemically modified peptides

| *protein* | *probe* | *captured peptide* | *modification* | *parent ion intensity* | *charge* | *m/z*$_{theor.}$ | *average m/z*$_{meas.}$ | *error (ppm)* | *SD* | *n* |
|---|---|---|---|---|---|---|---|---|---|---|
| SCRN3 | **PHZyne** | Glox*DTFVALPPATVGNR | $TEV_{heavy}$ | 5.0E+07 | 3 | 706.3810 | 706.3812 | 0.2 | 0.0008 | 96[†] |
| | | Glox*DTFVALPPATVGNR | $TEV_{light}$ | 4.0E+07 | 3 | 704.3764 | 704.3766 | 0.3 | 0.0010 | 123[†] |
| | | Pyvl*DTFVALPPATVGNR | $TEV_{heavy}$ | 3.5E+07 | 2 | 1066.0756 | 1066.0757 | 0.1 | 0.0007 | 248[†] |
| | | Pyvl*DTFVALPPATVGNR | $TEV_{light}$ | 2.5E+07 | 2 | 1063.0687 | 1063.0689 | 0.2 | 0.0007 | 234[†] |
| | **PHAyne** | Glox*DTFVALPPATVGNR | $TEV_{heavy}$ | 1.3E+08 | 2 | 1045.0521 | 1045.0522 | 0.1 | 0.0008 | 108[†] |
| | | Glox*DTFVALPPATVGNR | $TEV_{light}$ | 1.0E+08 | 2 | 1042.0452 | 1042.0454 | 0.2 | 0.0007 | 132[†] |
| | | Pyvl*DTFVALPPATVGNR | $TEV_{heavy}$ | 8.0E+07 | 2 | 1052.0600 | 1052.0599 | -0.1 | 0.0012 | 65[†] |
| | | Pyvl*DTFVALPPATVGNR | $TEV_{light}$ | 5.0E+07 | 2 | 1049.0531 | 1049.0532 | 0.1 | 0.0009 | 52[†] |

* site of modification by probe clicked with TEV tags.
[†] the sum of detections ($n \geq 52$) in MS1 spectra from two technical replicates.
Glox: glyoxylyl modification.
Pyvl: pyruvoyl modification.

### Table S2. Constructs of PHZyne mouse targets

| Construct (species) | source | identifier |
|---|---|---|
| MAOA_pCMV6 (mouse) | OriGene | MR208442 |
| MAOB_pCMV6 (mouse) | OriGene | MR226825 |
| MAOA_pRK5 (mouse) | This paper | |
| MAOB_pRK5 (mouse) | This paper | |
| PIPOX_pcDNA3.1+/C-(K)-DYK (mouse) | GenScript | clone ID OMu03158D |
| PIPOXC319A_pcDNA3.1+/C-(K)-DYK (mouse) | GenScript | clone ID OMu03158D |
| PCYOX1_pcDNA3.1+/C-(K)-DYK (mouse) | GenScript | clone ID OMu01091D |
| PCYOX1Y455A_pcDNA3.1+/C-(K)-DYK (mouse) | GenScript | clone ID OMu01091D |
| ALDH2_pcDNA3.1+/C-(K)-DYK (mouse) | GenScript | clone ID OMu18018D |
| ALDH2C321A_pcDNA3.1+/C-(K)-DYK (mouse) | GenScript | clone ID OMu18018D |
| SCRN3_ pcDNA3.1+/C-(K)-DYK (mouse) | GenScript | clone ID OMu06436D |
| SCRN2_ pcDNA3.1+/C-(K)-DYK (mouse) | GenScript | Clone ID OMu12145 |
| SCRN3_pRK5 (mouse) | This paper | |
| SCRN2_pRK5 (mouse) | This paper | |
| SCRN3C6A_pRK5 (mouse) | This paper | |
| SCRN2C12A_pRK5 (mouse) | This paper | |

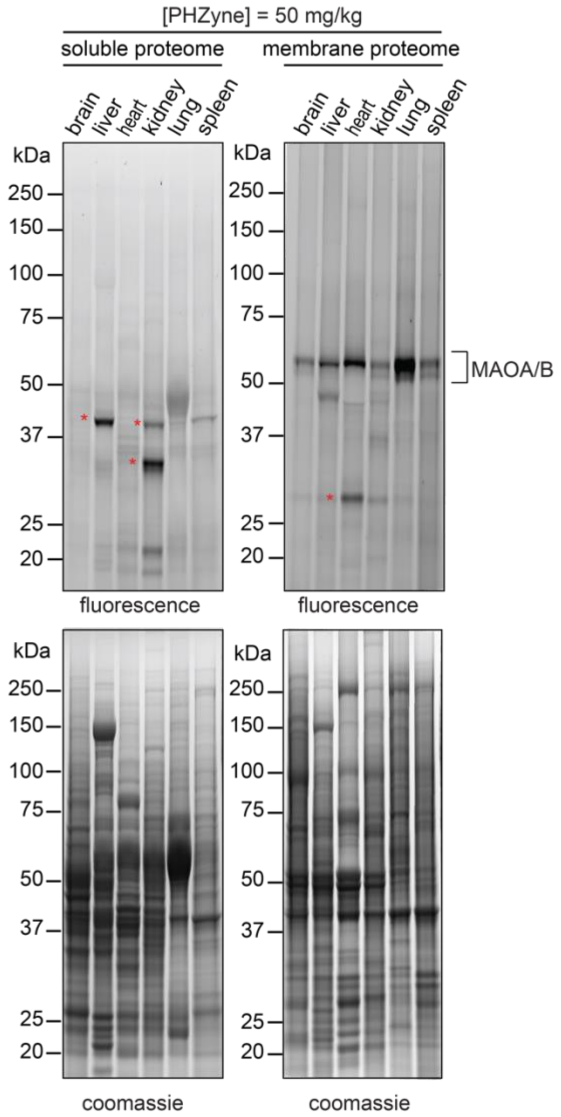


**Figure S1. Gel-based profiles for PHZyne-treated mouse tissues.** SDS-PAGE analysis of soluble and membrane proteomes from six mouse tissues (brain, liver, heart, kidney, lung, and spleen) harvested 4 h post-injection PHZyne (I.P., 50 mg/kg) and in-gel fluorescence scanning of PHZyne-modified proteins as measured by CuAAC to a rhodamine-azide tag (*upper*). Corresponding expression profiles are shown after Coomassie staining (*lower*). Doses and molecular weight markers are indicated.

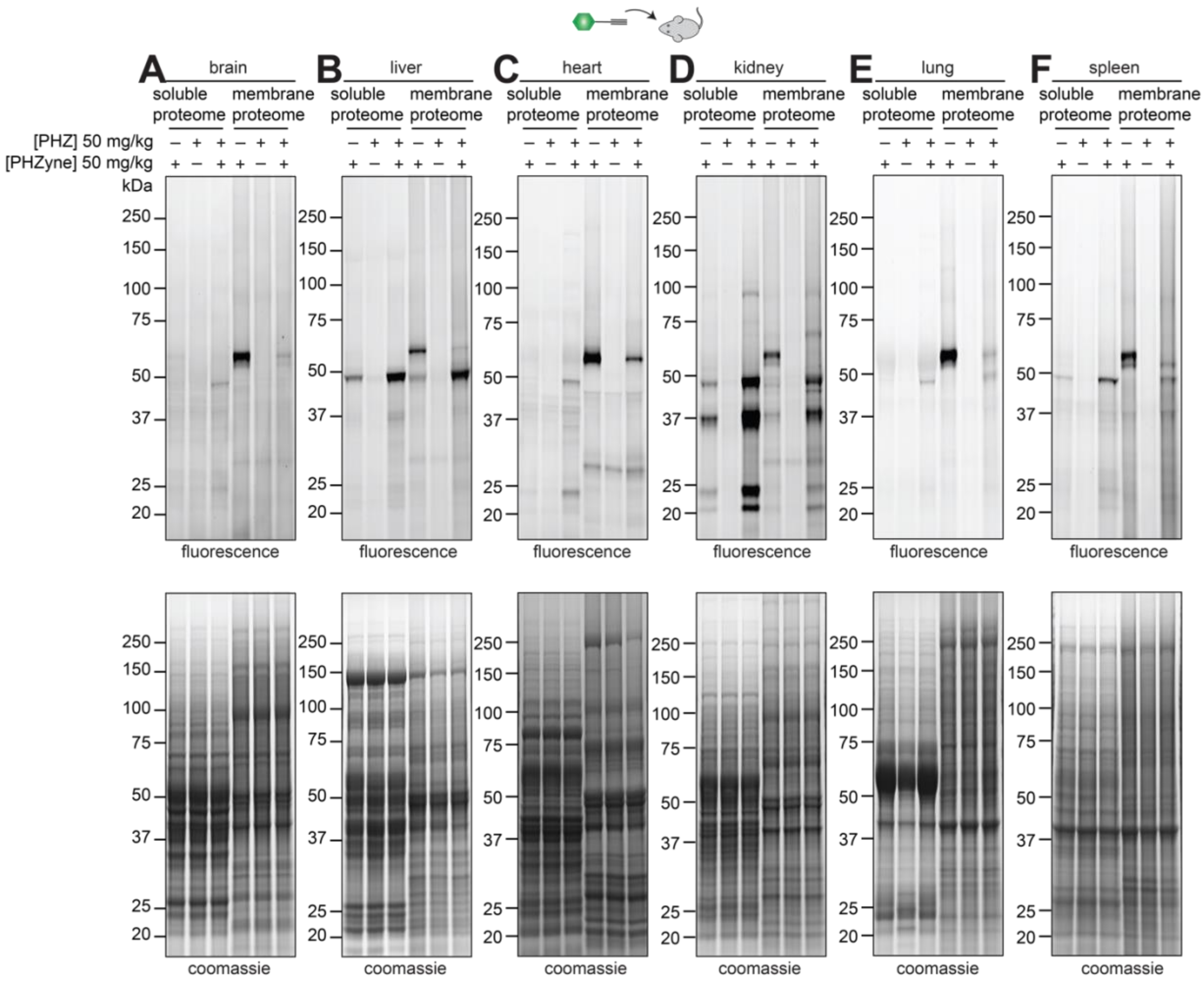


**Figure S2. Gel-based profiles for systematic PHZyne/PHZ-treated mouse tissues.** SDS-PAGE analysis of soluble and membrane proteomes from mouse **(A)** brain, **(B)** liver, **(C)** heart, **(D)** kidney, **(E)** lung, or **(F)** spleen tissues harvested 4 h post-injection with PHZyne and/or PHZ (I.P., 50 mg/kg) and in-gel fluorescence scanning (*upper*). Corresponding expression profiles are shown after Coomassie staining (*lower*). Doses and molecular weight markers are indicated.

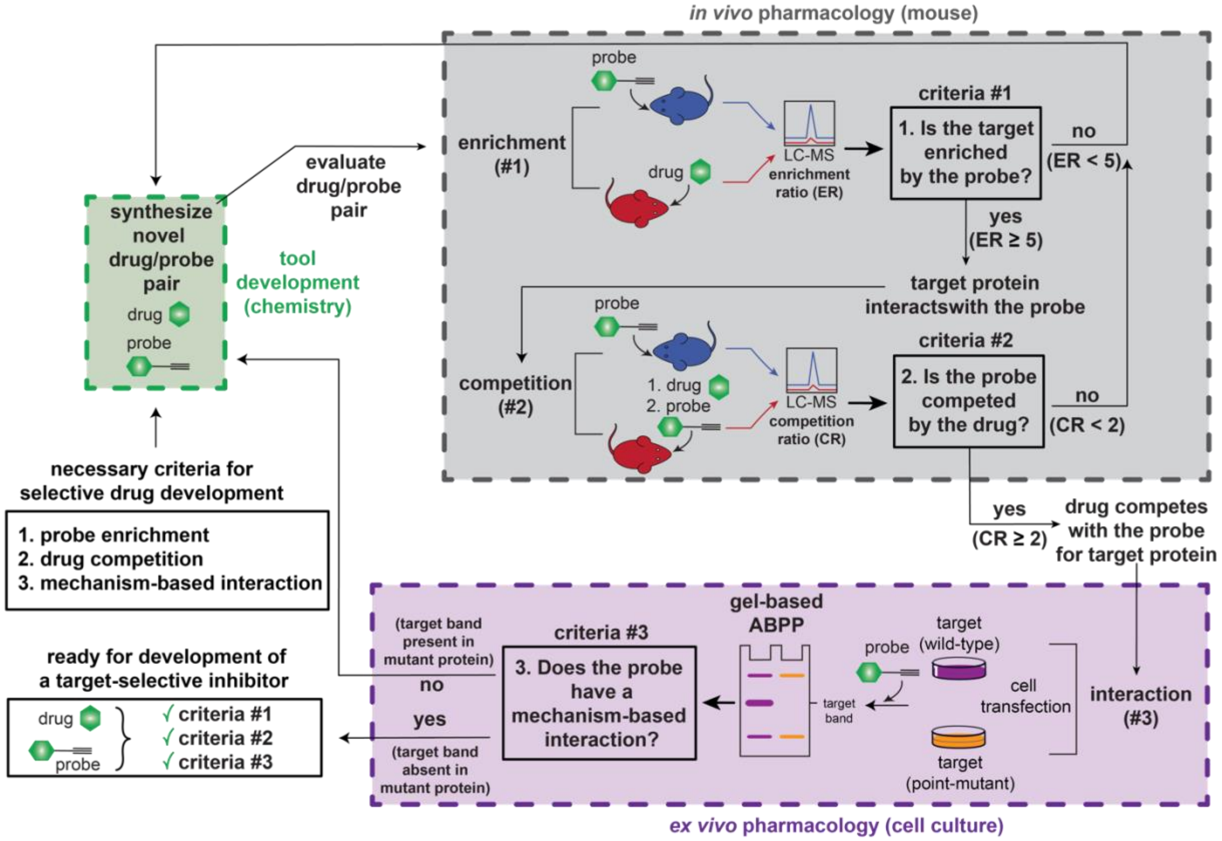


**Figure S3. Expanded schematic for target identification using a probe/drug pair**. MS-based proteomics experiments (enrichment and competition) identify and quantify probe/drug-target interactions *in vivo* that are reconstituted in cells to validate and characterize the site and mechanistic basis for the interactions captured *in vivo*

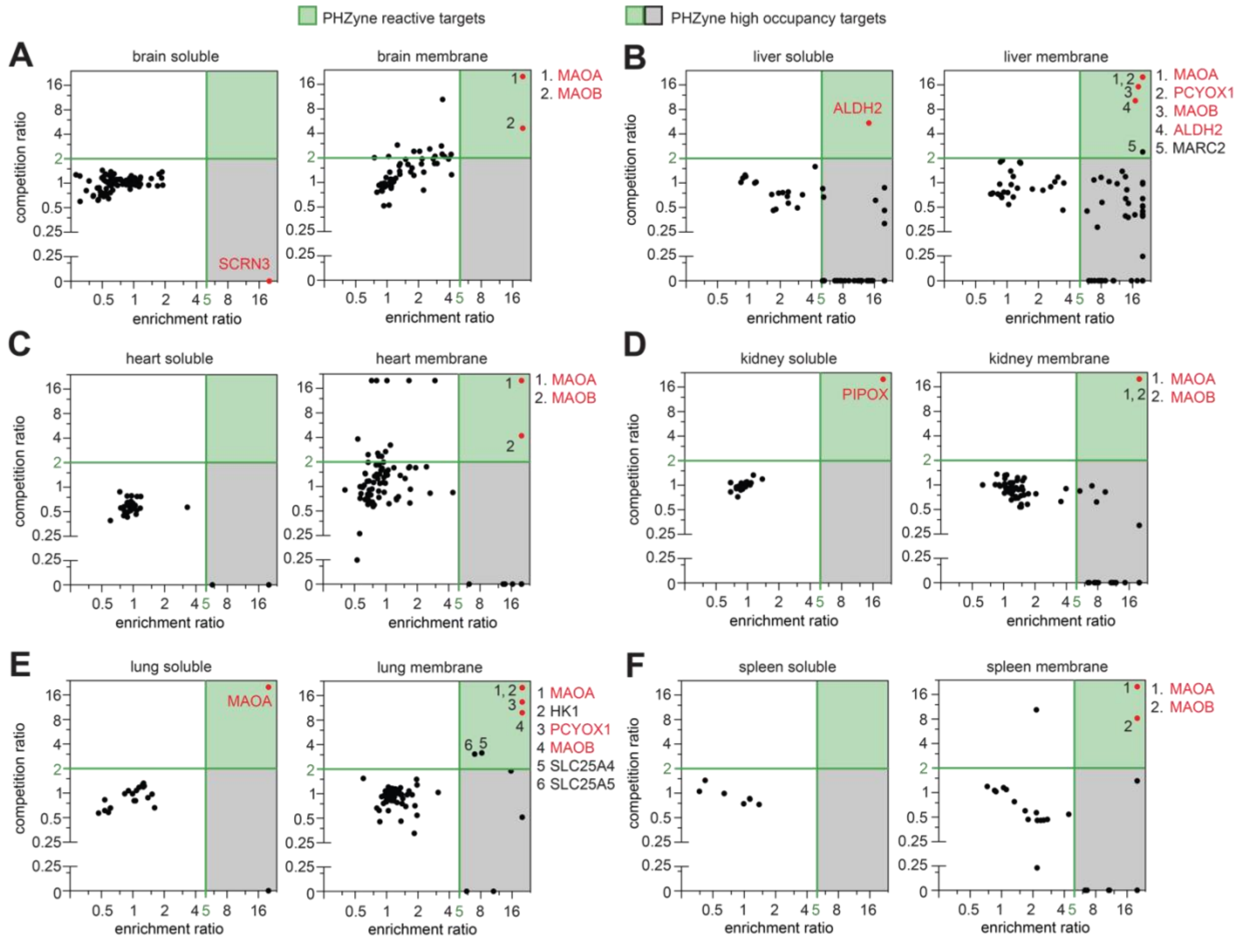


**Figure S4. Identification of PHZyne targets in mouse tissues.** Quadrant plot of average competition versus enrichment ReDiMe ratios for PHZyne/PHZ targets in the soluble (*left*) and membrane (*right*) proteomes of mouse **(A)** brain, **(B)** liver, **(C)** heart, **(D)** kidney, **(E)** lung, and **(F)** spleen from quantitative proteomic experiments ($n \geq 4$). Proteins with ratios that were quantified in enrichment experiments but were either undetected or did not pass the criteria for quantification in corresponding competition experiments were plotted along the x-axis. Proteins with ER ≥ 5 and CR ≥ 2 (upper right quadrant shown in green) were considered both high-reactivity and high-occupancy; listed to the right of the plot. Proteins shown in red are further investigated in this work.

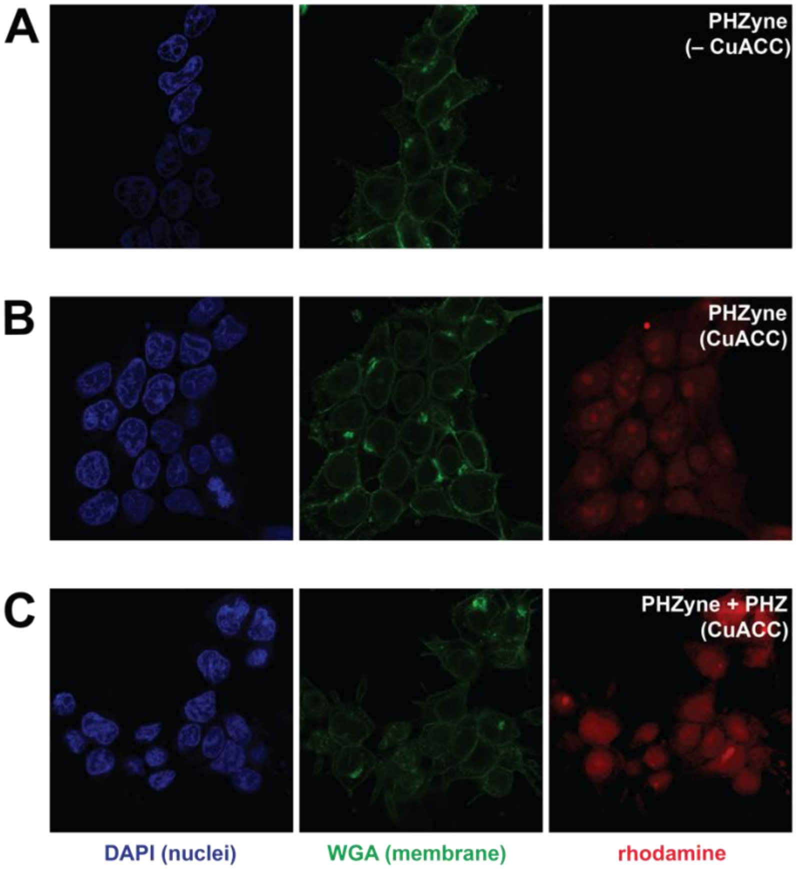


**Figure S5. Raw image data for PHZyne-reactivity in HEK293T cells**. PHZyne-labelling in the absence **(A)** and presence **(B)** of CuAAC and in competition with PHZ **(C).** Nuclei and membranes were stained with DAPI (blue, *left* panels*)* and WGA (green, *middle* panels), respectively**.** Rhodamine is conjugated to the probe by CuAAC (magenta, *right* panels). Merged image data and analysis shown in **Fig. 3**.

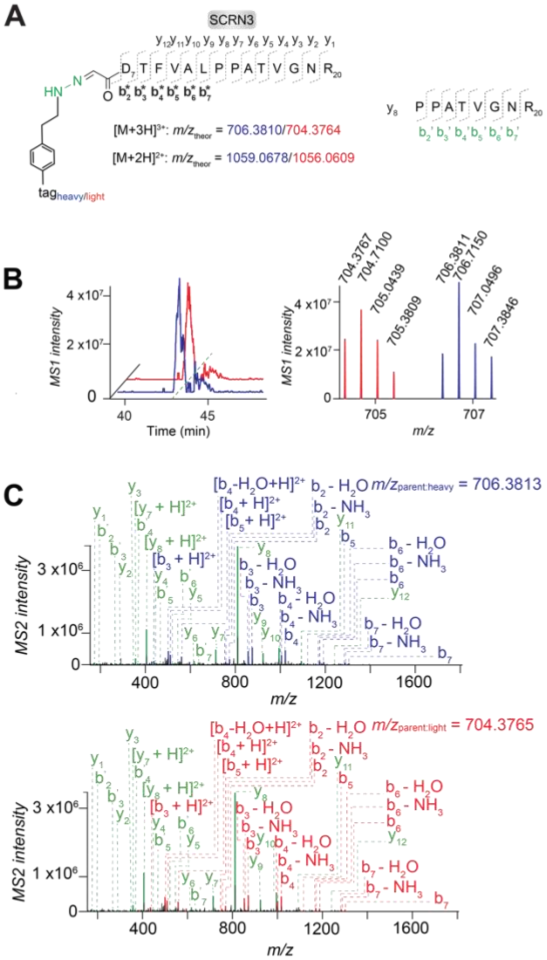


**D**

| | Theoretical $m/z$ | | Measured $m/z$ (error in ppm) | |
|---|---|---|---|---|
| | Heavy | Light | Heavy | Light |
| $b_2$ | 775.3878 | 769.3740 | 775.3888 (1.3) | 769.3743 (0.4) |
| $b_3$ | 876.4335 | 870.4217 | 876.4364 (3.3) | 870.4219 (0.2) |
| $b_4$ | 1023.5039 | 1017.4901 | 1023.5048 (0.9) | 1017.4903 (0.2) |
| $b_5$ | 1122.5723 | 1116.5585 | 1122.5745 (2.0) | 1116.5598 (1.2) |
| $b_6$ | 1193.6094 | 1187.5956 | 1193.6099 (0.4) | 1187.5933 (-1.9) |
| $b_7$ | 1306.6935 | 1300.6797 | 1306.6949 (1.1) | 1300.6753 (-3.4) |
| $[b_2 + H]^{2+}$ | 388.1976 | 385.1907 | 388.1981 (1.3) | 385.1908 (0.3) |
| $[b_3 + H]^{2+}$ | 438.7204 | 435.7145 | 438.7218 (3.2) | 435.7149 (0.9) |
| $[b_4 + H]^{2+}$ | 512.2556 | 509.2487 | 512.2560 (0.8) | 509.2491 (0.8) |
| $[b_5 + H]^{2+}$ | 561.7898 | 558.7829 | 561.7903 (0.9) | 558.7831 (0.4) |
| $[b_6 + H]^{2+}$ | 597.3084 | 594.3016 | 597.3093 (1.5) | 594.3007 (-1.5) |
| $b_2$-$NH_3$ | 758.3613 | 752.3475 | 758.3644 (4.1) | 752.3492 (2.3) |
| $b_3$-$NH_3$ | 859.4070 | 853.3952 | 859.4153 (9.7) | 853.4010 (6.8) |
| $b_4$-$NH_3$ | 1006.4774 | 1000.4636 | 1006.4795 (2.1) | 1000.4650 (1.4) |
| $b_5$-$NH_3$ | 1105.5458 | 1099.5320 | 1105.5497 (3.5) | 1099.5338 (1.6) |
| $b_6$-$NH_3$ | 1176.5829 | 1170.5691 | 1176.5847 (1.5) | 1170.5720 (2.5) |
| $b_7$-$NH_3$ | 1289.6670 | 1283.6532 | 1289.6700 (2.3) | 1283.6584 (4.1) |
| $b_2$-$H_2O$ | 757.3772 | 751.3634 | 757.3776 (0.5) | 751.4102 (62.3) |
| $b_3$-$H_2O$ | 858.4249 | 852.4111 | 858.4258 (1.0) | 852.4111 (0.0) |
| $b_4$-$H_2O$ | 1005.4933 | 999.4795 | 1005.4955 (2.2) | 999.4800 (0.5) |
| $b_5$-$H_2O$ | 1104.5617 | 1098.5479 | 1104.5636 (1.7) | - |
| $b_6$-$H_2O$ | 1175.5989 | 1169.5851 | 1175.5980 (-0.8) | 1169.5825 (-2.2) |
| $b_7$-$H_2O$ | 1288.6829 | 1282.6691 | 1288.6891 (4.8) | 1282.6658 (-2.6) |
| $[b_3$-$H_2O + H]^{2+}$ | 429.7161 | 426.7092 | 429.7168 (1.6) | 426.7095 (0.7) |
| $[b_4$-$H_2O + H]^{2+}$ | 503.2503 | 500.2434 | 503.2510 (1.4) | 500.2440 (1.2) |
| $[y_8 + H]^{2+}$ | 406.2247 | | 406.2251 (1.0) | 406.2249 (0.5) |
| $[y_7 + H]^{2+}$ | 357.6983 | | 357.6986 (0.8) | 357.6985 (0.6) |
| $y_{12}$ | 1241.7001 | | 1241.6855 (-11.8) | 1241.7028 (2.2) |
| $y_{11}$ | 1094.6317 | | 1094.6324 (0.6) | 1094.6312 (-0.5) |
| $y_{10}$ | 995.5633 | | 995.5637 (0.4) | 995.5632 (-0.1) |
| $y_9$ | 924.5261 | | 924.5270 (1.0) | 924.5261 (0.0) |
| $y_8$ | 811.4421 | | 811.4426 (0.6) | 811.4420 (-0.1) |
| $y_7$ | 714.3893 | | 714.3903 (1.4) | 714.3899 (0.8) |
| $y_6$ | 617.3365 | | 617.3375 (1.6) | 617.3376 (1.8) |
| $y_5$ | 546.2994 | | 546.2986 (-1.5) | 546.2996 (0.4) |
| $y_4$ | 445.2518 | | 445.2522 (0.9) | 445.2523 (1.1) |
| $y_3$ | 346.1833 | | 346.1839 (1.7) | 346.1835 (0.6) |
| $y_2$ | 289.1619 | | 289.1623 (1.4) | 289.1629 (3.5) |
| $y_1$ | 175.1190 | | 175.1190 (0.0) | 175.1190 (0.0) |
| $b'_7$ | 637.3304 | | 637.3320 (2.5) | 637.3284 (-3.1) |
| $b'_6$ | 523.2875 | | 523.2872 (-0.6) | 523.2874 (-0.2) |
| $b'_5$ | 466.2660 | | 466.2656 (-0.9) | 466.2659 (-0.2) |
| $b'_4$ | 367.1976 | | 367.1986 (2.7) | 367.1983 (1.9) |
| $b'_3$ | 266.1499 | | 266.1500 (0.4) | 266.1501 (0.8) |
| $b'_2$ | 195.1128 | | 195.1127 (-0.5) | 195.1131 (1.5) |

**Figure S6. MS characterization of PHZyne-labelled glyoxylyl in mouse SCRN3**. **(A)** structures and theoretical parent masses of heavy- and light-tagged SCRN3 peptides labelled by PHZyne and processed by the isoTOP-ABPP method. **(B)** Parent EICs and corresponding isotopic envelopes for heavy- (blue) and light- (red) tagged peptides detected from mouse SCRN3-transfected HEK293T cells. **(C)** MS2 spectra generated from parent ions. Unshifted ions are shown in green and the shifted ion series in blue (heavy) and red (light). **(D)** Summary table of theoretical versus observed spectra assignments generated under high-resolution MS2 conditions.

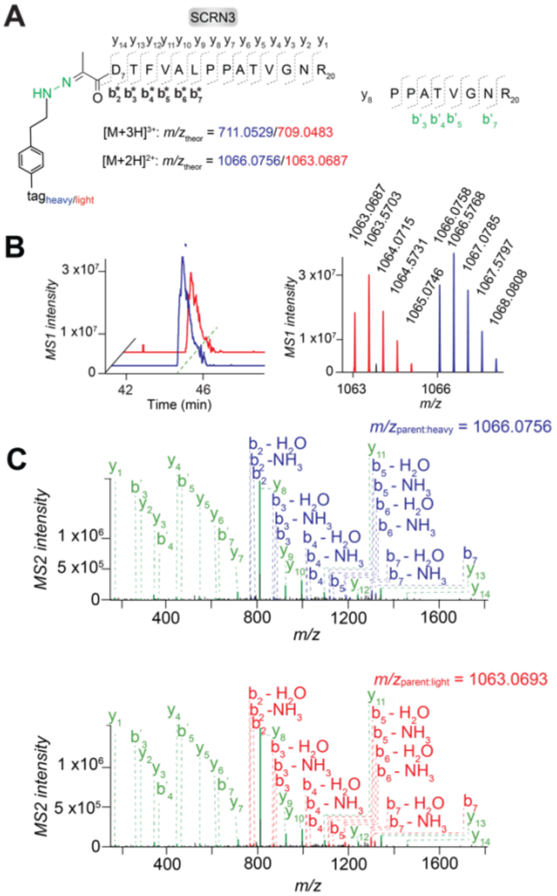


**D**

| | Theoretical $m/z$ Heavy | Theoretical $m/z$ Light | Measured $m/z$ (error in ppm) Heavy | Measured $m/z$ (error in ppm) Light |
|---|---|---|---|---|
| $b_2$ | 789.4035 | 783.3897 | 789.4047 (1.5) | 783.3888 (-1.1) |
| $b_3$ | 890.4512 | 884.4374 | 890.4544 (3.6) | 884.4388 (1.6) |
| $b_4$ | 1037.5196 | 1031.5058 | 1037.5220 (2.3) | 1031.5179 (11.7) |
| $b_5$ | 1136.5880 | 1130.5742 | 1136.5852 (-2.5) | 1130.5768 (2.3) |
| $b_6$ | 1207.6251 | 1201.6113 | 1207.6266 (1.2) | 1201.6088 (-2.1) |
| $b_7$ | 1320.7092 | 1314.6954 | 1320.7100 (0.6) | 1314.6947 -0.5) |
| $b_2$-$NH_3$ | 772.3770 | 766.3622 | 772.3793 (3.0) | 766.3652 (3.9) |
| $b_3$-$NH_3$ | 873.4247 | 867.4109 | 873.4357 (12.6) | 867.4115 (0.7) |
| $b_4$-$NH_3$ | 1020.4931 | 1014.4793 | 1020.4932 (0.1) | 1014.4819 (2.6) |
| $b_5$-$NH_3$ | 1119.5615 | 1113.5477 | 1119.5651 (3.2) | 1113.5529 (4.7) |
| $b_6$-$NH_3$ | 1190.5986 | 1184.5848 | 1190.6012 (2.2) | 1184.5880 (2.7) |
| $b_7$-$NH_3$ | 1303.6827 | 1297.6689 | 1303.6858 (2.4) | 1297.6718 (2.2) |
| $b_2$-$H_2O$ | 771.3929 | 765.3791 | 771.3923 (-0.8) | 765.3783 (-1.0) |
| $b_3$-$H_2O$ | 872.4406 | 866.4268 | 872.4414 (0.9) | 866.4233 (-4.0) |
| $b_4$-$H_2O$ | 1019.5090 | 1013.4952 | 1019.5090 (0.0) | 1013.4914 (-3.7) |
| $b_5$-$H_2O$ | 1118.5774 | 1112.5636 | 1118.57501 (-2.1) | 1112.5634 (-0.2) |
| $b_6$-$H_2O$ | 1189.6146 | 1183.6008 | 1189.6219 (6.1) | 1183.6044 (3.0) |
| $b_7$-$H_2O$ | 1302.6986 | 1296.6848 | 1302.6981 (-0.4) | 1296.6842 (-0.5) |
| $y_{14}$ | 1457.7747 | | 1457.7784 (2.5) | 1457.7755 (0.5) |
| $y_{13}$ | 1342.7478 | | 1342.7483 (0.4) | 1342.7472 (-0.4) |
| $y_{12}$ | 1241.7001 | | 1241.7000 (-0.1) | 1241.6962 (-3.1) |
| $y_{11}$ | 1094.6317 | | 1094.6315 (-0.2 | 1094.6317 (0.0) |
| $y_{10}$ | 995.5633 | | 995.5632 (-0.1) | 995.5631 (-0.2) |
| $y_9$ | 924.5261 | | 924.5263 (0.2) | 924.5261 (0.0) |
| $y_8$ | 811.4421 | | 811.4422 (0.1) | 811.4424 (0.4) |
| $y_7$ | 714.3893 | | 714.3897 (0.6) | 714.3900 (1.0) |
| $y_6$ | 617.3365 | | 617.3354 (-1.8) | 617.3384 (3.1) |
| $y_5$ | 546.2994 | | 546.2997 (0.5) | 546.3004 (1.8) |
| $y_4$ | 445.2518 | | 445.2518 (0.0) | 445.2521 (0.7) |
| $y_3$ | 346.1833 | | 346.1833 (0.0) | 346.1837 (1.2) |
| $y_2$ | 289.1619 | | 289.1620 (0.3) | 289.1613 (-2.1) |
| $y_1$ | 175.1190 | | 175.1187 (-1.7) | 175.1191 (0.6) |
| $b'_7$ | 637.3304 | | 637.3301 (-0.5) | 637.3316 (1.9) |
| $b'_5$ | 466.2660 | | 466.2658 (-0.4) | 466.2661 (0.2) |
| $b'_4$ | 367.1976 | | 367.1984 (2.2) | 367.1982 (1.6) |
| $b'_3$ | 266.1499 | | 266.1490 (-3.4) | 266.1504 (1.9) |

**Figure S7. MS characterization of PHZyne-labelled pyruvoyl in mouse SCRN3**. **(A)** structures and theoretical parent masses of heavy- and light-tagged SCRN3 peptides labelled by PHZyne and processed by the isoTOP-ABPP method. **(B)** Parent EICs and corresponding isotopic envelopes for heavy- (blue) and light- (red) tagged peptides detected from mouse SCRN3-transfected HEK293T cells. **(C)** MS2 spectra generated from parent ions. Unshifted ions are shown in green and the shifted ion series in blue (heavy) and red (light). **(D)** Summary table of theoretical versus observed spectra assignments generated under high-resolution MS2 conditions.

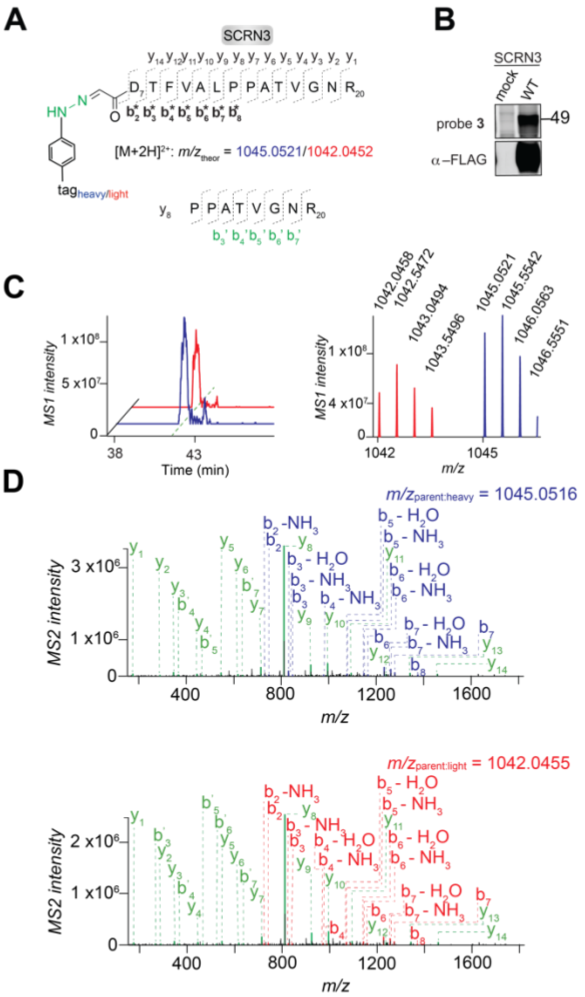


**E**

| | Theoretical $m/z$ Heavy | Theoretical $m/z$ Light | Measured $m/z$ (error in ppm) Heavy | Measured $m/z$ (error in ppm) Light |
|---|---|---|---|---|
| $b_2$ | 747.3565 | 741.3427 | 747.3558 (-0.9) | 741.3417 (-1.3) |
| $b_3$ | 848.4042 | 842.3904 | 848.4124 (9.7) | 842.3882 (-2.6) |
| $b_4$ | 995.4726 | 989.4588 | - | 989.4510 (-7.9) |
| $b_5$ | 1094.5410 | 1088.5272 | - | 1088.5237 (-3.2) |
| $b_6$ | 1165.5781 | 1159.5643 | 1165.5822 (3.5) | 1159.5621 (-1.9) |
| $b_7$ | 1278.6622 | 1272.6484 | 1278.6613 (-0.7) | 1272.6466 (-1.4) |
| $b_8$ | 1375.7150 | 1369.7012 | 1375.7124 (-1.9) | 1369.7073 (4.5) |
| $b_7$-$NH_3CO$ | 1233.6407 | 1227.6269 | 1233.6398 (-0.7) | 1227.6259 (-0.8) |
| $b_2$-$NH_3$ | 730.3300 | 724.3162 | 730.3309 (1.2) | 724.3134 (-3.9) |
| $b_3$-$NH_3$ | 831.3777 | 825.3639 | 831.3928 (18.2) | 825.3882 (29.4) |
| $b_4$-$NH_3$ | 978.4461 | 972.4323 | 978.4496 (3.6) | 972.4319 (-0.4) |
| $b_5$-$NH_3$ | 1077.5145 | 1071.5007 | 1077.5187 (3.9) | 1071.5103 (9.0) |
| $b_6$-$NH_3$ | 1148.5516 | 1142.5378 | 1148.5529 (1.1) | 1142.5382 (0.4) |
| $b_7$-$NH_3$ | 1261.6357 | 1255.6219 | 1261.6382 (2.0) | 1255.6223 (0.3) |
| $b_3$-$H_2O$ | 830.3936 | 824.3798 | 830.3945 (1.1) | - |
| $b_4$-$H_2O$ | 977.4620 | 971.4482 | - | 971.4520 (3.9) |
| $b_5$-$H_2O$ | 1076.5304 | 1070.5166 | 1076.5314 (0.9) | 1070.5143 (-2.1) |
| $b_6$-$H_2O$ | 1147.5676 | 1141.5538 | 1147.5815 (12.1) | 1141.5686 (13.0) |
| $b_7$-$H_2O$ | 1260.6516 | 1254.6378 | 1260.6510 (-0.5) | 1254.6350 (-2.2) |
| $y_{14}$ | 1457.7747 | | 1457.7708 (-2.7) | 1457.7725 (-1.5) |
| $y_{13}$ | 1342.7478 | | 1342.7476 (-0.1) | 1342.7444 (-2.5) |
| $y_{12}$ | 1241.7001 | | 1241.7015 (1.1) | 1241.6978 (-1.9) |
| $y_{11}$ | 1094.6317 | | 1094.6315 (-0.2) | 1094.6305 (-1.1) |
| $y_{10}$ | 995.5633 | | 995.5624 (-0.9) | 995.5625 (-0.8) |
| $y_9$ | 924.5261 | | 924.5253 (-0.9) | 924.5254 (-0.8) |
| $y_8$ | 811.4421 | | 811.4414 (-0.9) | 811.4418 (-0.4) |
| $y_7$ | 714.3893 | | 714.3892 (-0.1) | 714.3898 (0.7) |
| $y_6$ | 617.3365 | | 617.3375 (1.6) | 617.3369 (0.6) |
| $y_5$ | 546.2994 | | 546.2991 (-0.5) | 546.2999 (0.9) |
| $y_4$ | 445.2518 | | 445.2516 (-0.4) | 445.2519 (0.2) |
| $y_3$ | 346.1833 | | 346.1831 (-0.6) | 346.1838 (1.4) |
| $y_2$ | 289.1619 | | 289.1622 (1.0) | 289.1627 (2.8) |
| $y_1$ | 175.1190 | | 175.1186 (-2.3) | 175.1191 (0.6) |
| $b'_7$ | 637.3304 | | 637.3253 (-8.0) | 637.3317 (2.0) |
| $b'_6$ | 523.2875 | | - | 523.2878 (0.6) |
| $b'_5$ | 466.2660 | | 466.2659 (-0.2) | 466.2666 (1.3) |
| $b'_4$ | 367.1976 | | 367.1979 (0.8) | 367.1978 (0.5) |
| $b'_3$ | 266.1499 | | - | 266.1504 (1.9) |

**Figure S8. MS characterization of PHAyne-labelled glyoxylyl in mouse SCRN3**. **(A)** structures and theoretical parent masses of heavy- and light-tagged SCRN3 peptides labelled by PHAyne and processed by the isoTOP-ABPP method. **(B)** PHAyne-labelling of wild-type mouse SCRN3. Probe labelling (*upper*) and expression profiles (*lower*) for probe-treated cells expressing SCRN3. Transfection with the appropriate empty expression vector ('mock') is used as a control, molecular weights are indicated. **(C)** parent EICs and corresponding isotopic envelopes for heavy- (blue) and light- (red) tagged peptides detected from SCRN3-transfected HEK293T cells. **(D)** MS2 spectra generated from parent ions. Unshifted ions are shown in green and the shifted ion series in blue (heavy) and red (light). **(E)** Summary table of theoretical versus observed spectra assignments generated under high-resolution MS2 conditions.

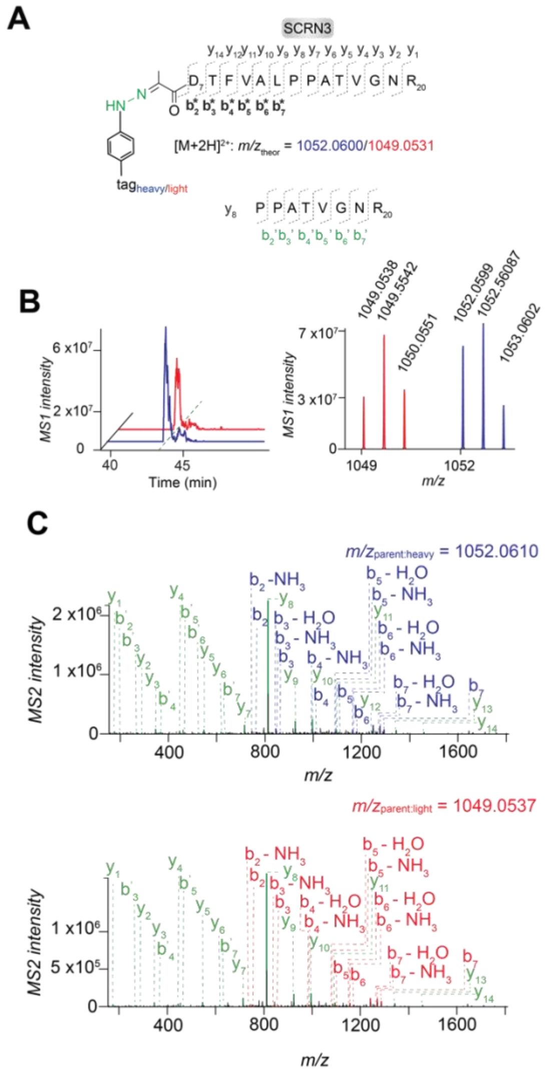


D

| | Theoretical *m/z* | | Measured *m/z* (error in ppm) | |
|---|---|---|---|---|
| | Heavy | Light | Heavy | Light |
| $b_2$ | 761.3722 | 755.3584 | 761.3711 (-1.4) | 755.3593 (1.2) |
| $b_3$ | 862.4199 | 856.4061 | 862.4266 (7.8) | 856.4164 (12.0) |
| $b_4$ | 1009.4883 | 1003.4745 | 1009.4974 (9.0) | - |
| $b_5$ | 1108.5567 | 1102.5429 | 1108.5580 (1.2) | 1102.5403 (-2.4) |
| $b_6$ | 1179.5938 | 1173.5800 | 1179.5963 (2.1) | 1173.5835 (3.0) |
| $b_7$ | 1292.6779 | 1286.6641 | 1292.6780 (0.1) | 1286.6700 (4.6) |
| $b_7$-$NH_3CO$ | 1247.6564 | 1241.6426 | 1247.6563 (-0.1) | 1241.6476 (4.0) |
| $b_2$-$NH_3$ | 744.3457 | 738.3319 | 744.3469 (1.6) | 738.3278 (-5.6) |
| $b_3$-$NH_3$ | 845.3934 | 839.3796 | 845.4073 (16.4) | 839.3784 (-1.4) |
| $b_4$-$NH_3$ | 992.4618 | 986.4480 | 992.4675 (5.7) | 986.4486 (0.6) |
| $b_5$-$NH_3$ | 1091.5302 | 1085.5164 | 1091.5331 (2.7) | 1085.5212 (4.4) |
| $b_6$-$NH_3$ | 1162.5673 | 1156.5535 | 1162.5723 (4.2) | 1156.5576 (3.5) |
| $b_7$-$NH_3$ | 1275.6514 | 1269.6376 | 1275.6539 (2.0) | 1269.6392 (1.3) |
| $b_3$-$H_2O$ | 844.4093 | 838.3955 | 844.4095 (0.2) | - |
| $b_4$-$H_2O$ | 991.4777 | 985.4639 | - | 985.4642 (0.3) |
| $b_5$-$H_2O$ | 1090.5461 | 1084.5323 | 1090.5547 (7.9) | 1084.5286 (-3.4) |
| $b_6$-$H_2O$ | 1161.5833 | 1155.5695 | 1161.5951 (10.2) | 1155.5782 (7.5) |
| $b_7$-$H_2O$ | 1274.663 | 1268.6535 | 1274.6672 (3.3) | 1268.6534 (-0.1) |
| $y_{14}$ | 1457.7747 | | 1457.7739 (-0.5) | 1457.7698 (3.4) |
| $y_{13}$ | 1342.7478 | | 1342.7474 (-0.3) | 1342.7457 (-1.6) |
| $y_{12}$ | 1241.7001 | | 1241.6997 (-0.3) | - |
| $y_{11}$ | 1094.6317 | | 1094.6311 (-0.5) | 1094.6315 (-0.2) |
| $y_{10}$ | 995.5633 | | 995.5626 (-0.7) | 995.5629 (-0.4) |
| $y_9$ | 924.5261 | | 924.5257 (-0.4) | 924.5259 (-0.2) |
| $y_8$ | 811.4421 | | 811.4417 (-0.5) | 811.4423 (0.2) |
| $y_7$ | 714.3893 | | 714.3893 (0.0) | 714.3902 (1.3) |
| $y_6$ | 617.3365 | | 617.3368 (0.5) | 617.3363 (-0.3) |
| $y_5$ | 546.2994 | | 546.2988 (-1.1) | 546.3002 (1.5) |
| $y_4$ | 445.2518 | | 445.2516 (-0.4) | 445.2522 (0.9) |
| $y_3$ | 346.1833 | | 346.1830 (-0.9) | 346.1838 (1.4) |
| $y_2$ | 289.1619 | | 289.1613 (-2.1) | 289.1624 (1.7) |
| $y_1$ | 175.1190 | | 175.1184 (-3.4) | 175.1190 (0.0) |
| $b'_7$ | 637.3304 | | 637.3309 (0.8) | 637.3299 (-0.8) |
| $b'_6$ | 523.2875 | | 523.2860 (-2.9) | - |
| $b'_5$ | 466.2660 | | 466.2656 (-0.9) | 466.2670 (2.1) |
| $b'_4$ | 367.1976 | | 367.1967 (-2.5) | 367.1973 (-0.8) |
| $b'_3$ | 266.1499 | | 266.1488 (-4.1) | 266.1503 (1.5) |
| $b'_2$ | 195.1128 | | 195.1126 (-1.0) | - |

**Figure S9. MS characterization of PHAyne-labelled pyruvoyl in mouse SCRN3**. **(A)** structures and theoretical parent masses of heavy- and light-tagged SCRN3 peptides labelled by PHAyne and processed by the isoTOP-ABPP method. **(B)** parent EICs and corresponding isotopic envelopes for heavy- (blue) and light- (red) tagged peptides detected from SCRN3-transfected HEK293T cells. **(C)** MS2 spectra generated from parent ions. Unshifted ions are shown in green and the shifted ion series in blue (heavy) and red (light). **(D)** Summary table of theoretical versus observed spectra assignments generated under high-resolution MS2 conditions.

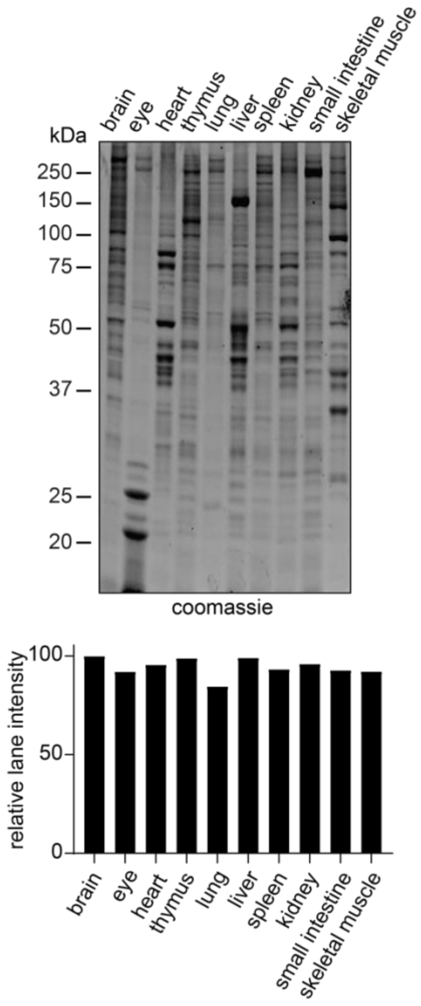


**Figure S10. Coomassie staining as loading control for SCRN3 protein expression in mouse tissues.** Expression profiles for mouse tissues after Coomassie staining (*upper*). Whole gel lane intensities, normalized relative to intensity in brain, are shown below.

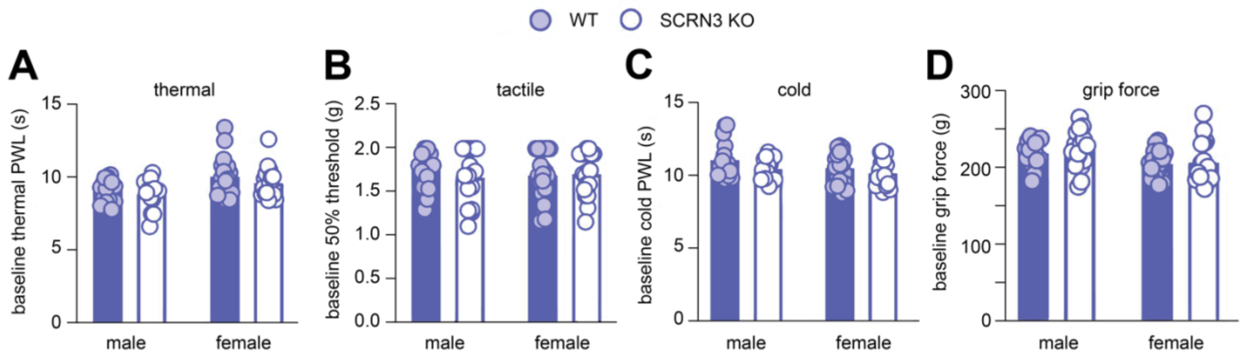


**Figure S11. Baseline measurements for four behavioral tests prior to IT injection.** Behavioral assessment of pain was measured in male and female WT and SCRN3 KO mice by **(A)** thermal hyperalgesia using hotplate assay, **(B)** tactile allodynia using von Frey test, **(C)** cold allodynia using cold plantar assay, and **(D)** paw grip strength deficit using digital grip force meter. Behavioral tests were analyzed using 2-way ANOVA (sex x genotype) and Bonferroni post hoc. Data are presented as mean ± SEM. No statistically significant differences were observed between WT and SCRN3 KO males and females.
